# Selective Sugar Transport Supports *Proteus mirabilis* Fitness in the Urinary Tract

**DOI:** 10.64898/2026.02.20.707088

**Authors:** Allyson E. Shea, Shiuhyang Kuo, Surbhi Gupta, Sara N. Smith, Trishna Appaji, Taylor Mitchell, Harry L.T. Mobley, Melanie M. Pearson

## Abstract

*Proteus mirabilis* is a leading cause of complicated urinary tract infections (UTIs). Prior work showed *P. mirabilis* metabolizes sugars during experimental UTI, yet the role of sugar import systems in pathogenesis remains poorly defined. To investigate this, we generated a panel of 47 targeted mutants in predicted sugar transporter genes and assessed their growth *in vitro* and fitness *in vivo*. Growth screening in nutrient-rich and minimal media revealed carbon source-dependent defects in several phosphotransferase system (PTS) mutants, including *ptsH* and *ptsI*. Pooled insertion sequencing (In-seq) identified *xapB*, *ptsH*, and *ptsI* as *in vivo* fitness factors, with validation in a traditional murine co-challenge model. Functional studies showed that *xapB*, annotated as a xanthosine permease, did not support xanthosine or guanosine uptake in *P. mirabilis*, suggesting misannotation. Dissection of the PTS network revealed that a triple mutant lacking *scrA*, *ulaC*, and *ptsG* recapitulated the *ptsH* phenotype *in vivo*. To evaluate whether increased sugar availability exacerbates these defects, we modeled glucosuria using the SGLT2 inhibitor dapagliflozin in CBA/J mice. Dapagliflozin treatment significantly increased urinary glucose and enhanced *P. mirabilis* colonization. There was an inverse correlation between colonization and urinary glucose, but only in untreated mice. These findings reveal limitations in genome-based transporter annotation, establish a functional link between sugar import and *P. mirabilis* fitness during UTI, and demonstrate that host metabolic conditions such as glucosuria can influence the severity of infection.

**AUTHOR SUMMARY:** All living organisms require nutrients to grow, survive, and cause disease. Bacteria like *Proteus mirabilis*, which causes urinary tract infections, rely on specialized systems to import and metabolize sugars available in the host environment. In this study, we systematically disrupted 47 genes predicted to encode sugar transporters in *P. mirabilis* and tested their contribution to infection in a mouse model. We identified three key genes (*xapB*, *ptsH*, and *ptsI*) that were critical for colonization. Further analysis showed that many sugar transporters in *P. mirabilis* were misannotated, and predicted substrates like sucrose and cellobiose were not utilized by the bacterium. We also demonstrated that high sugar conditions, mimicking diabetic urine using the drug dapagliflozin, worsened infection and increased disease severity. These results highlight the importance of carbohydrate acquisition for *P. mirabilis* during infection and emphasize the need to experimentally validate gene function rather than rely on predictions based on other bacteria like *E. coli*.

## INTRODUCTION

Urinary tract infections (UTIs) are among the most common bacterial infections, disproportionately affecting women and generating a substantial financial burden on the healthcare system (1, 2). In catheterized patients, UTIs are often polymicrobial and more likely to involve urease-producing species such as *Proteus mirabilis* (3, 4). This organism poses a particular clinical challenge in long-term catheterization, where its ability to hydrolyze urea raises urine pH, leading to the precipitation of magnesium ammonium phosphate (struvite) and calcium phosphate into crystalline deposits (5, 6). These deposits form kidney and bladder stones and obstruct catheters, contributing to recurrent infection, inflammation, and tissue damage (7, 8). Although well-studied in terms of urease activity, swarming motility, and fimbrial adhesion, the metabolic strategies used by *P. mirabilis* during infection remain incompletely understood.

Amino acids are the major carbon source available to microbes in the urinary tract and can be detected in gram quantities in the total daily urine output from healthy adults (9, 10). It is then logical that uropathogens would shift their resources to acquiring amino acids as a food source. This is indeed the case for uropathogenic *Escherichia coli* (UPEC), the major agent of uncomplicated UTI, which not only upregulates amino acid importers (11), but also possess greater redundancy in those systems compared to non-urinary isolates (12). This is in contrast to commensal *E. coli* in the gastrointestinal tract, which relies more heavily on mono- and disaccharides for growth (13). These same carbohydrates in the gut are in relatively low abundance in the urine.

Perhaps counterintuitively given this nutrient landscape, *P. mirabilis* activates a variety of metabolic and nutrient acquisition pathways during experimental UTI, including amino acid uptake, peptide transport, key enzymes of central metabolism, and, in particular, sugar uptake and catabolism (14). Specifically, the loss of glycolytic enzymes and the oxidative pentose phosphate pathway significantly reduce *P. mirabilis* colonization and fitness *in vivo* (15, 16). Genes in both the non-oxidative pentose phosphate pathway and phosphoglycerate kinase (*pgk*) have been deemed possible essential genes (16). This is in contrast to UPEC, where glycolytic genes are not induced during UTI and the loss of glycolysis had no effect; instead, gluconeogenesis is exclusively required (15, 17, 18). In fact, pathways involved in sugar import and catabolism were downregulated in UPEC collected directly from women with active UTI, while amino acid and peptide importers were upregulated (11). Collectively, this suggests that *P. mirabilis* has access to carbohydrate sources either in urine or from host cells within the urinary tract that confer an advantage during polymicrobial infection.

Diabetes is one prevalent health condition that results in an increase of available sugars in urine. Diabetic patients are at increased risk for contracting UTI (19) and a larger proportion of infections are caused by *P. mirabilis* (20). Sodium-glucose cotransporter 2 (SGLT2) is a common biological target to prevent the re-uptake of urinary glucose in diseases like diabetes, but also chronic kidney disease and heart failure (21). Use of SGLT2 inhibitors, such as dapagliflozin or canagliflozin, has been shown to increase bacterial burden in some murine models of UTI (22, 23). In particular, UPEC loads were significantly higher in the bladders and kidneys of dapagliflozin-treated mice, with elevated rates of dissemination to the spleen and liver observed at 48 hours post-inoculation (22). These findings suggest that elevated urinary glucose may worsen infection outcomes. We hypothesized that this effect would be more pronounced for *P. mirabilis*, which exhibits a greater dependence on glucose-driven metabolism during UTI.

To understand how *P. mirabilis* imports sugars during infection, we focused our study on sugar uptake systems. The best characterized family of these is the phosphoenolpyruvate (PEP) phosphotransferase system (PTS). The PTS comprises upstream regulatory proteins, including Enzyme I (EI) and the phosphocarrier protein HPr, which coordinate phosphorylation cascades that activate a wide array of sugar-specific importers (24). The recipients of this phosphorylation are the EII units, which coordinate to facilitate the uptake of one substrate or a small group of similarly related substrates (25, 26). In addition to importing sugars and sugar derivatives, this system has regulatory crosstalk related to nitrogen metabolism (27, 28), chemotaxis (29, 30), and virulence in some pathogens (25). In *P. mirabilis*, the identity of certain EII components and the corresponding predicted substrates relies heavily on annotations from other Gram-negative bacterial species. Given this, coupled with the known requirement for glycolysis during *P. mirabilis* colonization and virulence *in vivo*, we sought to better define the specific carbon sources fueling uropathogenicity.

Toward this goal, we generated a set of 47 mutants targeting predicted sugar importers and tested them in a murine model of UTI. Three mutants, including a putative xanthosine permease (*xapB*) and PTS core components *ptsH* and *ptsI*, exhibited significant *in vivo* fitness defects. A combination of three PTS substrate-specific mutations was sufficient to phenocopy the *ptsH* mutant *in vivo*, confirming that these systems collectively contribute to PTS-dependent fitness. Finally, using an SGLT2 inhibitor to induce hyperglucosuria in mice, we observed elevated bacterial burden and increased morbidity during experimental UTI. We also identified an inverse correlation between bacterial burden and urinary glucose in untreated mice that was abolished during hyperglucosuria from SGLT2 inhibition. Collectively, our data show that specific sugar uptake systems are required for *P. mirabilis* fitness during UTI, and glucosuria alters experimental UTI outcomes.

## RESULTS

### Generation and growth assessment of sugar transporter mutants in *Proteus mirabilis*

Glycolysis and related carbohydrate pathways have been shown to play a central role in *P. mirabilis* pathogenesis during urinary tract infection (UTI) (14, 15). To further define the contribution of sugar import and metabolism to bacterial fitness, we constructed a panel of targeted mutants in genes predicted to encode sugar transporters. Candidate genes were selected using multiple complementary approaches. First, phosphotransferase system (PTS) components were prioritized due to their known roles in carbohydrate uptake. In parallel, bioinformatic tools including KEGG (31), Transporter Classification Database (TCDB 2.0) (32), NCBI BLAST against sugar transporters encoded by UPEC CFT073 (33), NCBI Conserved Domains Database (34), Phyre2 (35), and the Bacterial and Viral Bioinformatics Resource Center (BV-BRC) (36) were used to identify additional sugar transport genes from ones annotated in the *P. mirabilis* HI4320 genome (37). To cast a wide net, we used a broad definition of sugar such that a prediction for sugar transport in any one database merited inclusion. This yielded a final list of 50 targets (**Table 1**). Mutants were generated using the targetron system for site-specific group II intron insertion (38, 39). Of the 50 genes targeted, we successfully created 47 mutants, each confirmed by PCR.

**Table 1.** Candidate P. mirabilis sugar transporter genes.

| ORF | NCBI | Name | Annotation<sup>a</sup> | Mutant | Type | Pool |
| --- | --- | --- | --- | --- | --- | --- |
| PMI0090 | PMI_RS00430 | *rbsB* | D-ribose ABC transporter, substrate-binding periplasmic protein | Y | ABC | 1 |
| PMI0091 | PMI_RS00435 | *rbsC* | ribose ABC transporter, permease protein | Y | ABC | 1 |
| PMI0092 | PMI_RS00440 | *rbsA* | ribose ABC transporter, ATP-binding protein | Y | ABC | 1 |
| PMI0093 | PMI_RS00445 | *rbsD* | high affinity ribose transport protein | Y | ABC | 1 |
| PMI0164 | PMI_RS00795 |  | MFS-family transporter | Y | MFS | 1 |
| PMI0291 | PMI_RS01405 | *treB* | PTS system, trehalose-specific IIBC component | Y | PTS | 2 |
| PMI0455 | PMI_RS02280 | *nagE* | pts system, N-acetylglucosamine-specific EIICBA component | Y | PTS | 2 |
| PMI0775 | PMI_RS03810 | *uup* | ABC transporter ATP-binding protein | Y | ABC | 1 |
| PMI0850 | PMI_RS04170 |  | MFS-family transporter | Y | MFS | 1 |
| PMI1175 | PMI_RS05670 | *fetB* | putative membrane protein | Y | ABC | 1 |
| PMI1176 | PMI_RS05675 |  | ABC transporter, ATP-binding protein | Y | ABC | 1 |
| PMI1224 | PMI_RS05905 |  | putative ABC-type multidrug transport system | Y | ABC | 1 |
| PMI1251 | PMI_RS06040 |  | MFS-family transporter | N | MFS | n/a |
| PMI1254 | PMI_RS06055 |  | MFS-family transporter | Y | MFS | 1 |
| PMI1515 | PMI_RS07345 | *sotB* | sugar efflux transporter | Y | MFS | 1 |
| PMI1570 | PMI_RS07650 | *xapB* | xanthosine permease (MFS-family transporter) | Y | MFS | 1 |
| PMI1775 | PMI_RS08705 | *ulaA* | putative ascorbate PTS system IIC component | Y | PTS | 2 |
| PMI1776 | PMI_RS08710 | *ulaB* | PTS system IIB component | Y | PTS | 2<sup>b</sup> |
| PMI1777 | PMI_RS08715 | *ulaC* | PTS system IIA component | Y | PTS | 2 |
| PMI1828 | PMI_RS09020 | *ptsH* | PTS system phosphocarrier protein | Y | PTS | 2 |
| PMI1829 | PMI_RS09025 | *ptsI* | phosphoenolpyruvate-protein phosphotransferase | Y | PTS | 2 |
| PMI1830 | PMI_RS09030 | *crr* | PTS family enzyme IIA component/glucose-specific phosphotransferase enzyme IIA component | Y | PTS | 2 |
| PMI1935 | PMI_RS09535 |  | L-lactate permease | Y | Others | 2 |
| PMI1943 | PMI_RS09575 |  | MFS-family transporter | N | MFS | n/a |
| PMI1946 | PMI_RS09590 | *sglT* | sodium/glucose cotransporter | Y | Others | 2 |
| PMI2101 | PMI_RS10350 | *bcsC* | cellulose synthase protein | Y | Others | 2 |
| PMI2135 | PMI_RS10520 | *agaF* | putative N-acetylgalactosamine-specific PTS system, IIA component | Y | PTS | 2 |
| PMI2136 | PMI_RS10525 | *agaD* | putative N-acetylgalactosamine-specific PTS system, IID component | Y | PTS | 2 |
| PMI2137 | PMI_RS10530 | *agaW* | putative N-acetylgalactosamine-specific PTS system, IIC component | Y | PTS | 2 |
| PMI2138 | PMI_RS10535 | *agaV* | putative N-acetylgalactosamine-specific PTS system, IIB component | Y | PTS | 2 |
| PMI2157 | PMI_RS10630 | *fruK* | 1-phosphofructokinase | Y | PTS | 2 |
| PMI2198 | PMI_RS10830 |  | MFS-family transporter | Y | MFS | 1 |
| PMI2226 | PMI_RS10970 | *bglF* | PTS system, IIBC component/PTS system beta-glucoside-specific IIA component, Glc family | Y | PTS | 2 |
| PMI2292 | PMI_RS11320 | *ptsG* | glucose-specific PTS system, EIIBC component | Y | PTS | 2 |
| PMI2597 | PMI_RS12825 | *nrpX* | MFS-family transporter | Y | MFS | 1 |
| PMI2649 | PMI_RS13050 |  | MFS-family transporter | Y | MFS | 1 |
| PMI2954 | PMI_RS14605 | *chbA* | N,N'-diacetylchitobiose-specific PTS system, IIA component | Y | PTS | 2 |
| PMI2955 | PMI_RS14610 | *chbC* | N,N'- diacetylchitobiose-specific PTS system, EIIC component | Y | PTS | 2 |
| PMI2956 | PMI_RS14615 | *chbB* | N,N'- diacetylchitobiose-specific PTS system, EIIB component | Y | PTS | 2 |
| PMI2967 | PMI_RS14670 |  | ABC-type sugar transport system, periplasmic component | Y | ABC | 1 |
| PMI2969 | PMI_RS14680 |  | MFS-family transporter | Y | MFS | 1 |
| PMI2982 | PMI_RS14745 |  | PTS system, EIIBC component | Y | PTS | 2 |
| PMI3099 | PMI_RS15340 |  | putative cellulose biosynthesis protein C | Y | Others | 2 |
| PMI3515 | PMI_RS17475 | *scrA* | putative sucrose PTS system IIBC component | Y | PTS | 2 |
| PMI3614 | PMI_RS17985 | *ugpC* | sn-glycerol-3-phosphate ABC transporter, ATP-binding protein | Y | ABC | 1 |
| PMI3615 | PMI_RS17990 | *ugpE* | sn-glycerol-3-phosphate ABC transporter, permease protein | Y | ABC | 1 |
| PMI3616 | PMI_RS17995 | *ugpA* | sn-glycerol-3-phosphate ABC transporter, permease protein | Y | ABC | 1 |
| PMI3617 | PMI_RS18000 | *ugpB* | glycerol-3-phosphate ABC transporter, substrate-binding protein | Y | ABC | 1 |
| PMI3646 | PMI_RS18140 | *ptsN* | nitrogen regulatory protein [includes: PTS system EIIA component] | N | PTS | n/a |
| PMI3705 | PMI_RS18435 |  | putative ABC transporter ATP-binding protein | Y | ABC | 1 |

<sup>a</sup>Annotations are from original Sanger Centre predictions or from KEGG
<sup>b</sup>PMI1776 was inadvertently left out of pool 2, and PMI1176 was included in both pools

We examined the growth of all 47 sugar transporter mutants in LB and in Minimal A medium supplemented with either 0.2% glucose or 0.2% glycerol as the sole carbon source (**Fig. 1**). Doubling time during logarithmic phase was calculated and compared to wild-type HI4320 (**Fig. 1A**). As expected, all mutants exhibited robust growth in LB, indicating that disruption of sugar-specific transport systems did not impair viability under nutrient-rich conditions. In contrast, four mutants displayed a statistically significant increase in doubling time when cultured in Minimal A with glucose as the carbon source (**Fig. 1B**), and all of them were in PTS-related genes. The *ptsG* mutant, encoding the EIIBC component of the glucose-specific PTS system (40), showed reduced growth (doubling time 52.63 min *vs.* 36.22 min for wt), consistent with its role in glucose import. Similarly, mutants lacking *crr* (53.41 min) and *ptsI* (51.23 min), which encode the carbohydrate repression resistance protein and Enzyme I of the PTS (41), respectively, were attenuated under these conditions. Notably, the *ptsH* mutant, which lacks the general phosphocarrier protein HPr, exhibited impaired growth in both glucose- (57.75 min) and glycerol- (49.28 min *vs.* 39.15 min for wt) containing Minimal A medium (**Fig. 1B-C**). We also observed a modest decrease in growth parameters for the *ptsI* mutant in glycerol (**Fig. 1C**), but this was not statistically significant (47.08 min). These findings align with the functional hierarchy of the PTS, as PtsI and PtsH comprise the phosphorelay for multiple sugar import systems and serve as central nodes for carbohydrate uptake and regulation in *P. mirabilis*, while Crr is paired with several substrate-specific importers including PtsG.

**Fig. 1.**
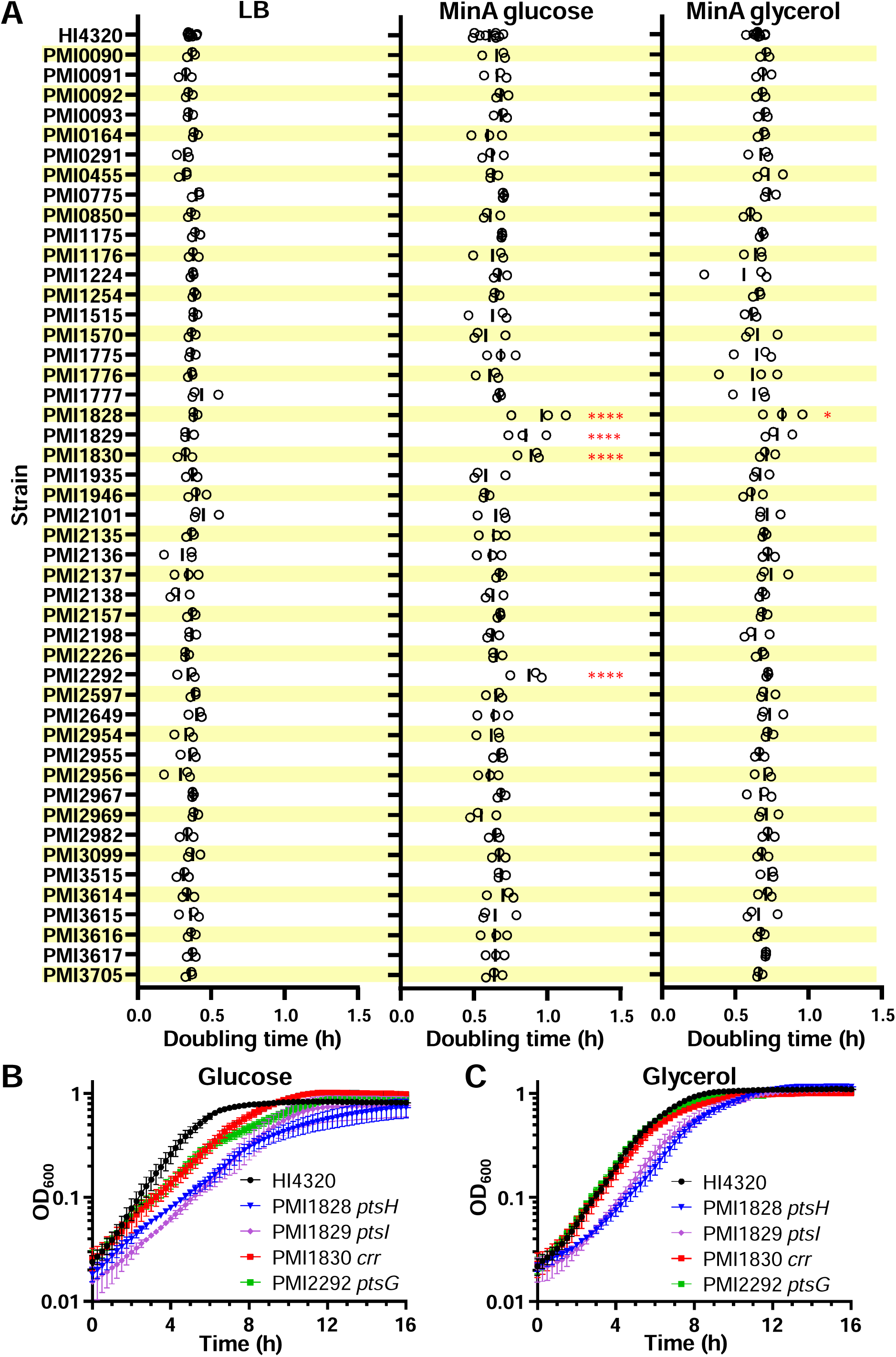
Growth parameters of 47 sugar transporter mutants. **A,** Doubling time measurements. Left, LB; middle, Minimal A with 0.2% glucose as the carbon source; right, Minimal A with 0.2% glycerol as the carbon source. Vertical lines show means. \**P*<0.05, \*\*\*\**P*<0.0001; one-way ANOVA *vs.* wild type HI4320 with Dunnett’s multiple comparisons test. **B-C,** Growth curves for PTS mutants with growth defects in Minimal A. **B,** Minimal A with 0.2% glucose as the carbon source. **C,** Minimal A with 0.2% glycerol as the carbon source. Error bars = SEM (n≥3).

### *In vivo* screening identifies *xapB*, *ptsH*, and *ptsI* as fitness factors

Previous work established that the bottleneck for *P. mirabilis* HI4320 in the murine model of ascending UTI limits reliable input pools to approximately 25 mutants (42). Based on this constraint, we divided our panel of 47 sugar metabolism mutants into two pools for *in vivo* testing (**Table 1**). Pool 1 consisted of 24 mutants from the ATP-binding cassette (ABC) and major facilitator superfamily (MFS) transporter families. Pool 2 included 23 mutants from the phosphotransferase system (PTS) and other single-gene predicted sugar transporters (**Fig. 2A**). Each pool was independently inoculated into the bladders of female CBA/J mice, and infection was allowed to proceed for 24 hours. Urine, bladder, and kidney samples were then harvested and individually barcoded for downstream sequencing (**Table S1**). The abundance of each mutant in the inoculum and recovered organs was determined using an insertion sequencing (In-seq) strategy (**Datasets S1 and S2**). An aliquot of each pool was plated to quantify the total input CFU and was also sequenced to identify variation that could have occurred from plating and outgrowth; no notable variation was observed in these control samples.

**Fig. 2.**
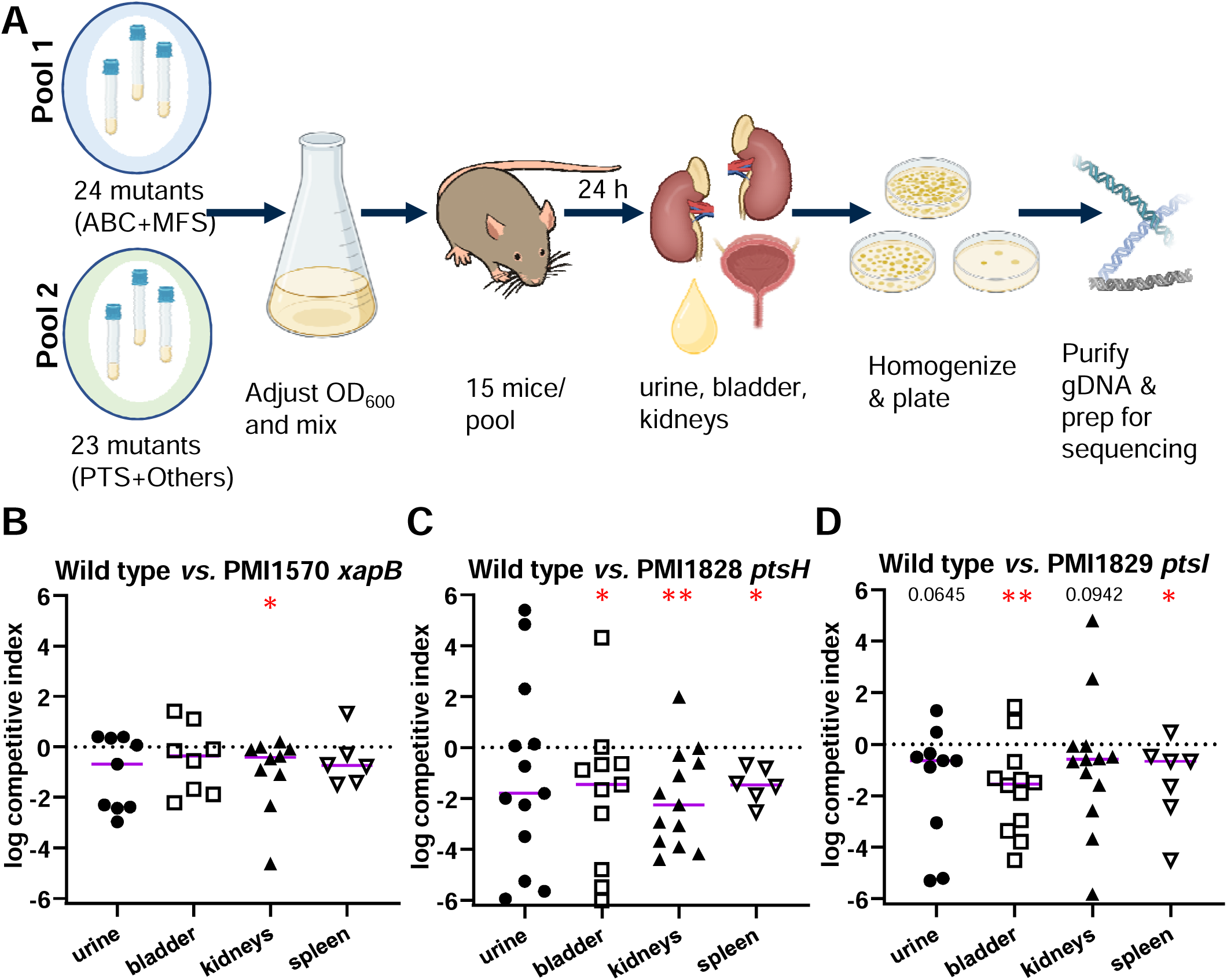
Identification of sugar importers that contribute to fitness during UTI. **A,** Experimental workflow for In-seq screening *P. mirabilis* sugar transporter mutants in mice. **B-D,** 1:1 wild type:mutant co-challenge competitive indices (7 dpi) for mutants identified in the In-seq screen. B, wild type *vs. xapB* (n = 10); **C,** wild type *vs. ptsH* (n = 15); **D,** wild type *vs. ptsI* (n = 15). Horizontal lines show medians. Dashed line indicates equal fitness of wild type and mutant (log CI= 0). \**P* < 0.05; \*\**P* < 0.01; exact value given for 0.10 > *P* > 0.05, one sample Wilcoxon test *vs.* a theoretical median of 0.

From Pool 1, the *xapB* mutant emerged as the only strain with a statistically significant fitness defect (**Fig. S1**). This mutant was underrepresented exclusively in the kidneys, suggesting an organ-specific role (**Fig. S1C**). *xapB* encodes a putative xanthosine permease of the MFS family, based on homology to the *E. coli* XapB transporter as predicted by KEGG (43, 44). From Pool 2, both *ptsH* and *ptsI* mutants were depleted in the bladder, and *ptsH* also showed a defect in the kidneys (**Fig. S2**). To validate the *in vivo* In-seq findings, each mutant (*xapB*, *ptsH*, and *ptsI*) was individually assessed in the established 7-day murine co-challenge model (45). For each strain, mice were inoculated transurethrally with a 1:1 mixture of mutant and wild-type HI4320, and competitive indices (CI) were calculated for urine, bladder, kidneys, and, to gauge dissemination into the bloodstream, spleens. The *xapB* mutant displayed a significant colonization defect in the kidneys (median log_10_ CI −0.4177) (**Figs. 2B, S3A**), consistent with the original In-seq data. The *ptsH* mutant exhibited significant defects in the bladder, kidney, and spleen (median log_10_ CI −1.459, −2.261, and −1.469, respectively), while *ptsI* was significantly outcompeted in the bladder and spleen (median log_10_ CI −1.538 and −0.6682, respectively) (**Figs. 2C-D, S3B-C**). Collectively, these experiments screened 47 sugar transport mutants for *in vivo* fitness and identified organ-specific defects that were reproducible in the traditional ascending UTI murine model.

### Functional characterization of putative xanthosine transporter XapB

To investigate the function of *xapB* and the other In-seq hits (*ptsH* and *ptsI*), we performed targeted phenotypic analyses guided by predicted functions of each transporter. We first focused on PMI1570, the putative xanthosine permease (*xapB*). When Minimal A medium was used with 0.1% xanthosine as the sole carbon source, HI4320 failed to grow (**Fig. 3A**). We therefore screened growth across Biolog Phenotype MicroArray carbon and nitrogen source plates to identify additional potential substrates in an unbiased fashion. No growth was observed when xanthosine was provided as a sole nitrogen source after 20 hours of incubation, indicating that strain HI4320 was unable to utilize xanthosine in this context (**Fig. 3B**). These findings suggest that XapB may not function as a xanthosine importer in *P. mirabilis* under the tested conditions. However, ruling xanthosine out as a carbon or nitrogen source does not exclude possibility for import.

**Fig. 3.**
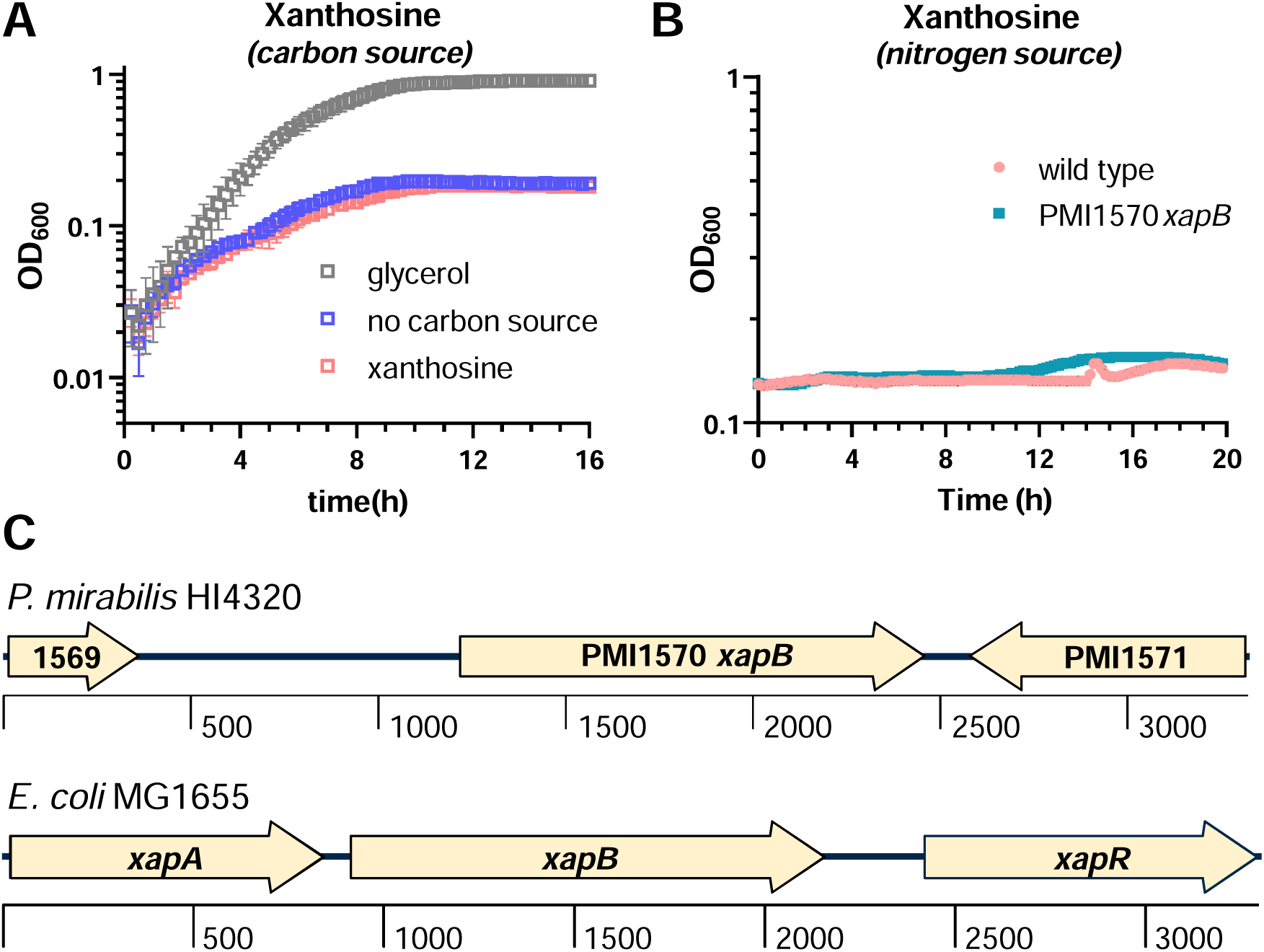
PMI1570 is likely misannotated as a xanthosine transporter. **A,** Growth curves for wild-type HI4320 in Minimal A with no carbon source, 0.2% glycerol, or 0.1% xanthosine as a sole carbon source (n = 3; error bars show SD). **B,** Growth on xanthosine as sole nitrogen source (Biolog plate PM3). **C,** The genetic organization of the putative *xap* locus in *P. mirabilis* (top) is missing elements from the *xap* locus in *E. coli*, where it has been characterized.

Further analysis of the genomic context revealed notable differences in operon structure compared to the *E. coli* MG1655 *xap* locus (**Fig. 3C**). In *P. mirabilis*, both *xapA*, encoding a xanthosine phosphorylase (PNP-II), and the regulatory gene *xapR* are absent. This single-gene organization raises the likelihood that XapB transports an alternate substrate. Beyond KEGG and BV-BRC’s annotations as a xanthosine transporter, other databases predict *xapB* to encode an undefined nucleoside transporter (BLAST and BioCyc). While *P. mirabilis* HI4320 XapB is 49% identical and 70% similar to *E. coli* MG1655 XapB, it is also 44% identical and 65% similar to the nucleoside:H^+^ symporter NupG. Biolog data suggested differential growth for wild-type HI4320 and the *xapB* mutant on guanosine (**Fig. S4 A-B**); however, the increase in optical density appeared to be due to a chemical precipitate, and no CFU were recovered at the experimental endpoint. There were no other notable differences in growth across 251 unique substrates between wild type and *xapB*. Nevertheless, we hypothesized that guanosine might enter through XapB.

Our group previously reported that a *guaA* mutant, which is deficient in GMP biosynthesis, exhibited impaired growth on murine organ agar, a severe defect *in vivo*, altered swarming motility, and a nutritional defect that could be rescued by exogenous RNA (42). As predicted, HI4320 was unable to use guanosine as either a sole nitrogen or carbon source in Minimal A medium (**Fig. 4A, B**). Despite this, exogenous guanosine successfully complemented the growth defect of the *guaA* mutant in Minimal A with glycerol, confirming the presence of a functional guanosine uptake system (**Fig. 4C**). A modest slowing of growth in the presence of guanosine was not due to the DMSO solvent, nor could this slowing be mitigated by using a lower concentration of guanosine (**Fig. S4 C-D**). To determine whether XapB was responsible for this activity, we constructed a *guaA*/*xapB* double mutant. As with the single *guaA* mutant, growth defects were rescued by guanosine supplementation (**Fig. 4D**). These data indicate that guanosine can enter the cell independent of XapB. The actual substrate of XapB, which must be available in the urinary tract and contributes to *P. mirabilis* fitness, remains to be determined.

**Fig. 4.**
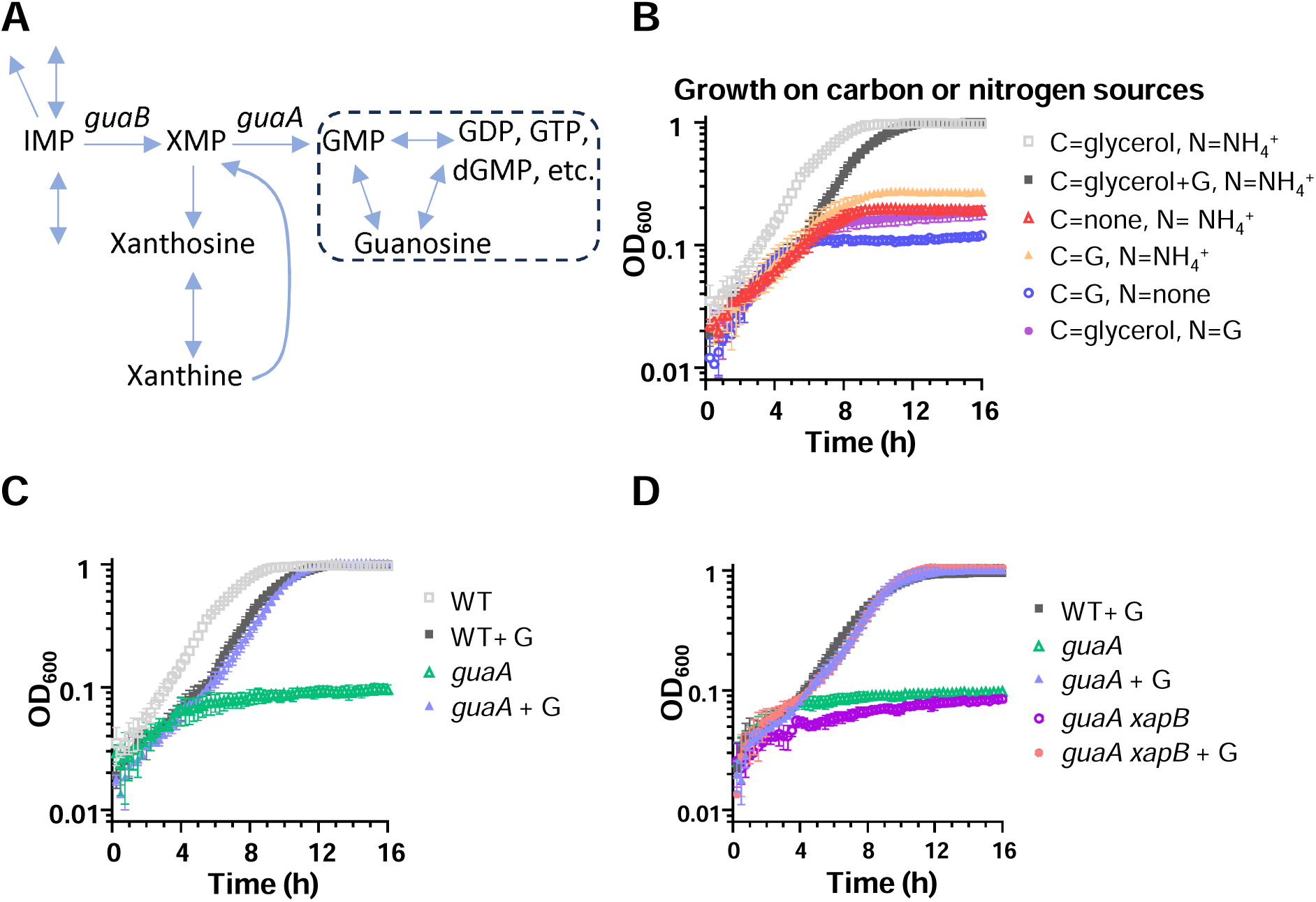
PMI1570 is not likely to be a guanosine transporter. **A,** Prediction of biochemical pathways encoded by *P. mirabilis* HI4320 including xanthosine and guanosine. *P mirabilis* is not predicted to be able to convert guanosine or related molecules into central metabolism (denoted by dashed outline). **B,** Growth curves in Minimal A indicated that wild-type HI4320 can’t use guanosine (G, 0.25 mg/mL) as a sole carbon (C) or nitrogen (N) source. **C-D,** Growth curves in Minimal A containing 0.2% glycerol. **C,** HI4320 can import guanosine to chemically complement the *guaA* mutant (wild type data are the same as in B). **D,** A *guaA xapB* double mutant can still be chemically complemented by guanosine, indicating that *xapB* probably doesn’t transport guanosine. In these experiments, guanosine was added at a final concentration of 0.05 mg/mL. For sole nitrogen source growth curves, n = 2. For all other conditions, n = 3. Error bars show SD.

### Dissecting the role of PTS components in *P. mirabilis* urinary tract fitness

The phosphotransferase system (PTS) is essential for carbohydrate uptake and consists of a relay of proteins that transfer phosphate from phosphoenolpyruvate (PEP) to sugar-specific permeases. In *P. mirabilis*, the PTS system appears to be organized similar to other Enterobacterales: *ptsI* encodes Enzyme I, which initiates the phospho-transfer cascade, and *ptsH* encodes HPr, a central phosphocarrier that relays phosphate to multiple specific import systems (46). *P. mirabilis* HI4320 is predicted to encode nine of these substrate-specific PTS importers (**Fig. 5**). Our *in vivo* co-challenge studies showed that both *ptsI* and *ptsH* single mutants were significantly outcompeted by wild type (**Fig. 2**). These broad defects suggested that loss of general PTS function impairs fitness, but the specific transporter systems, and thus substrates, responsible remained unclear. Of the nine importers, only glucose transporter *ptsG* had been experimentally confirmed for *P. mirabilis* HI4320 (**Fig. 1B**). To identify other PTS substrates that could contribute to the *ptsH* and *ptsI* fitness defects, we assessed growth of wild-type HI4320 and the *ptsH* mutant on 190 carbon sources and 95 nitrogen sources.

**Fig. 5.**
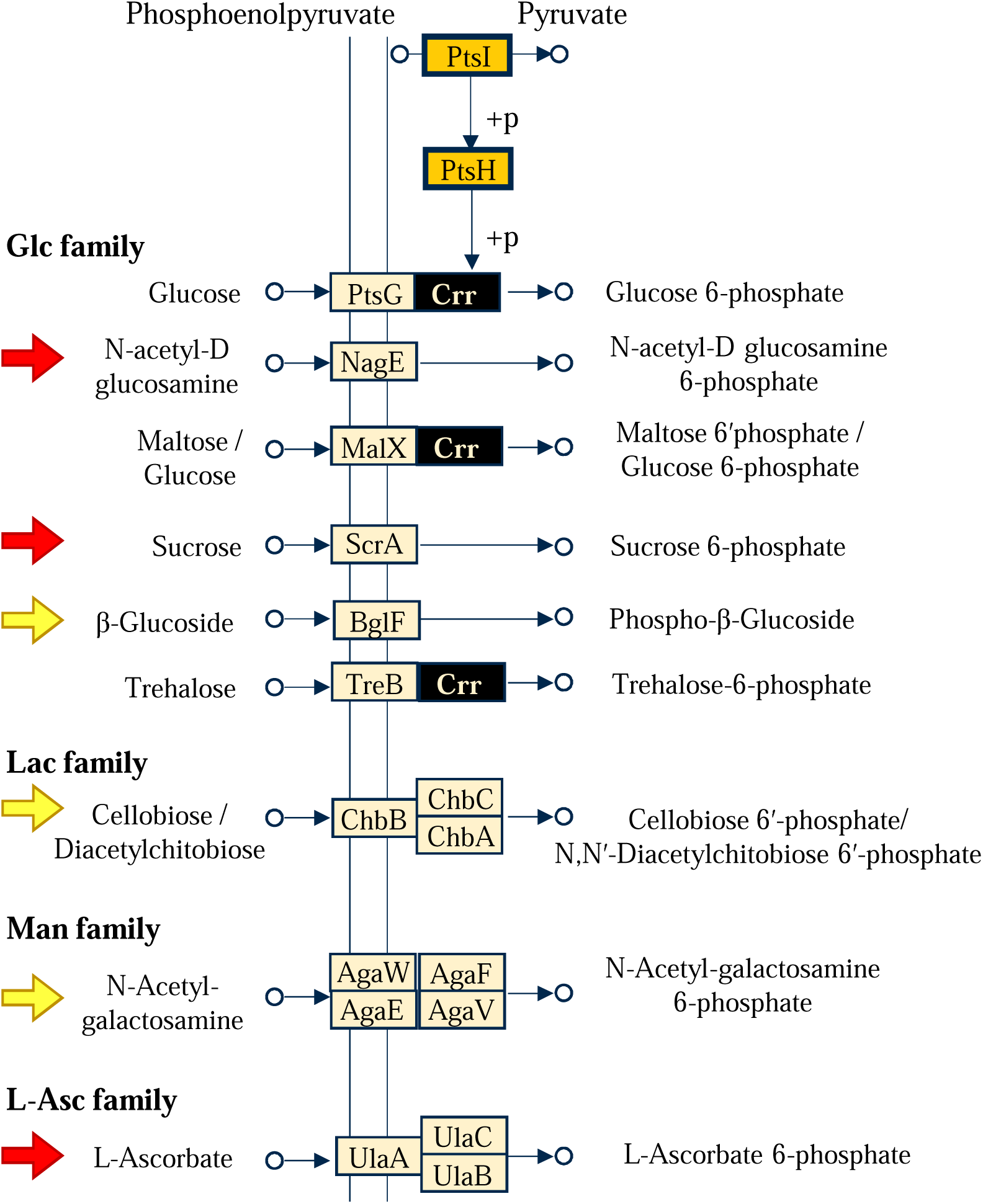
KEGG predictions for PTS-family transporters in *P. mirabilis* HI4320. Nine substrate-specific importers, shown here spanning the cell membrane depicted as parallel lines, are predicted to rely on the PtsHI phosphorelay. Crr (PTS enzyme IIA component) is predicted to interact with three substrate-specific importers and is highlighted in black. Arrows indicate two sets of non-*crr*-dependent triple mutants (red and yellow) that were tested in mice.

In addition to confirming glucose as a PTS substrate, we found the *ptsH* mutant had, as expected, greatly diminished growth on N-acetyl-D-glucosamine, N-acetyl-D-galactosamine, and trehalose (**Fig. 6 A-D, Table S1**). Predicted substrates of sucrose, maltose, and D-cellobiose were unable to support the growth of wild-type HI4320 (**Fig. 6 E-G**), suggesting possible misannotation. D-galactose did facilitate robust growth, but no differences were observed between wild type and mutant (**Fig. 6H**), which is consistent with the prediction that galactose is not a PTS substrate for *P. mirabilis*. Finally, the *ptsH* mutant displayed a growth defect, compared to wild type, in the presence of chondroitin sulfate C (**Fig. 6I**). This is interesting because chondroitin sulfate C is not a predicted PTS substrate, but a known chondroitinase in *P. mirabilis* likely facilitates cleavage of the substrate into glucuronic acid and PTS substrate N-acetylgalactosamine (47, 48). Collectively, we confirmed four predicted PTS substrates and did not identify any additional unpredicted PTS substrates that supported growth of *P. mirabilis* HI4320 (**Table S2**).

**Fig. 6.**
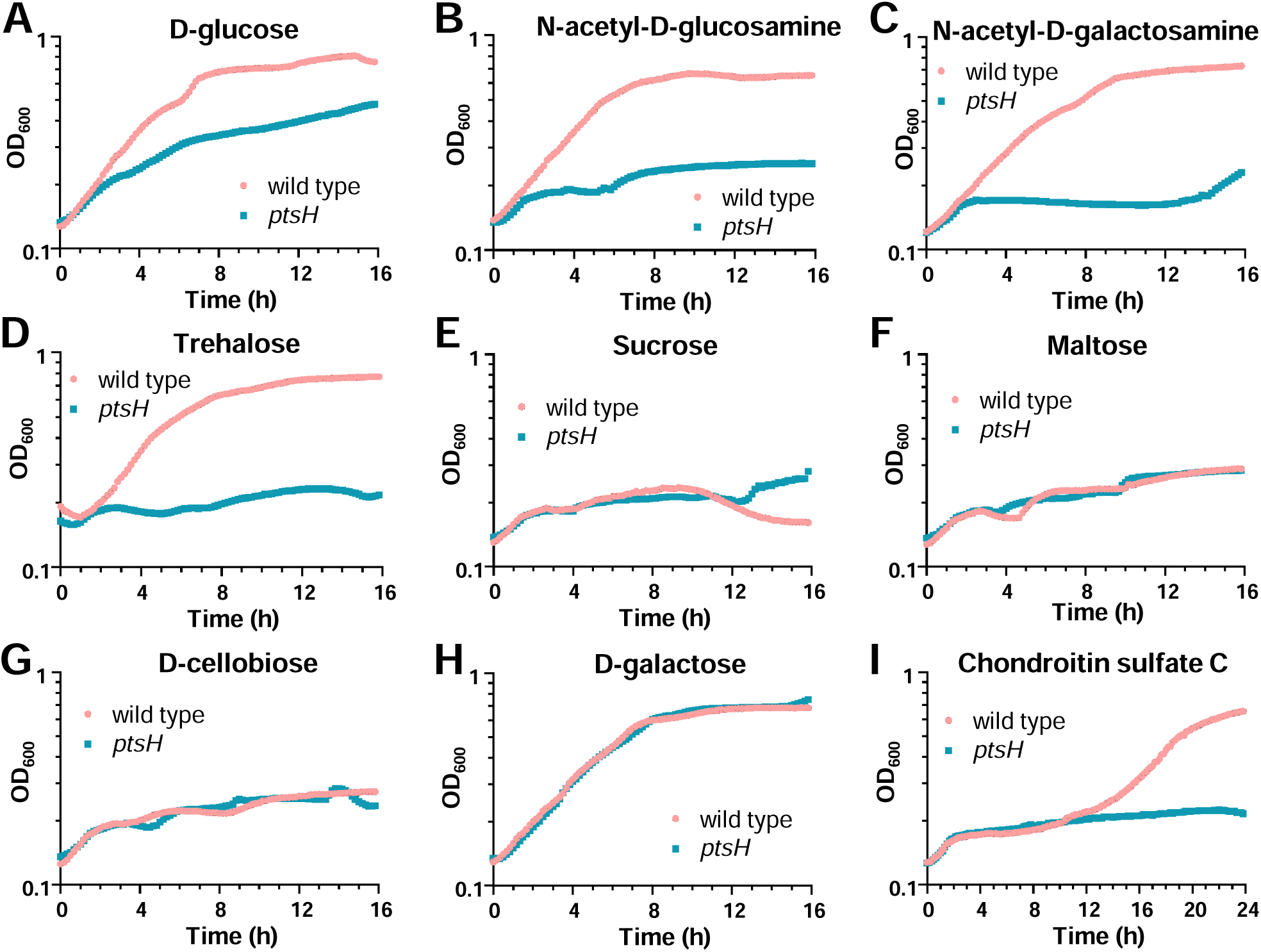
Notable carbon source growth curve results for wild type *vs.* the *ptsH* mutant. **A-H,** Predicted PTS substrates. **A-D,** Predicted PTS substrates that showed expected growth defects for the *ptsH* mutant. **E-G,** Predicted PTS substrates that did not support growth by wild-type HI4320, suggesting that their transporters might be misannotated. **H,** *P. mirabilis* utilizes galactose but it is not a PTS substrate. **I,** Chondroitin sulfate C was not predicted as a PTS substrate.

Remarkably, we detected growth defects of the *ptsH* mutant on various nitrogen sources (**Fig. S5**). Both ammonia and urea are preferred nitrogen sources that are particularly relevant to the urinary tract environment, and we observed a large growth defect of the *ptsH* mutant, compared to wild type, for both (**Fig. S5 A-B**). Similarly, L-amino acids such as leucine, arginine, histidine, phenylalanine, and cysteine also resulted in growth defects for the *ptsH* mutant (**Fig. S5 C-G**). Interestingly, nitrogen sources of serine and methionine did not yield differences between strains (**Fig. S5 H-I**). The nitrogen-dependent phenotypes we observed correlate with the growth defects of the *ptsH* and *ptsI* mutants in Minimal A with glycerol as the carbon source and provide an explanation why *ptsH* and *ptsI* had more severe defects with glucose as the carbon source compared with *ptsG* or *crr* mutants (**Fig. 1B-C**).

*P. mirabilis* swarming motility is induced by amino acid cues (49, 50). In addition, swarming defects often correlate with fitness defects during experimental UTI (51). Furthermore, a previous study in *Bacillus cereus* reported swarming defects for a *ptsH* mutant (52). For these reasons, we tested swarming motility and found that the *ptsH* mutant exhibited a modest but statistically significant reduction in swarming motility compared to wild-type HI4320 (**Fig. S6**).

To narrow down the specific substrate sugar import pathways that contribute to UTI fitness, we first excluded systems associated with Crr (PMI1830), an Enzyme IIA component not identified as a fitness factor in our In-seq screen (**Fig. S2, Fig. 5**). We then grouped the remaining PTS systems into categories of high or low/no induction during UTI based on previous *in vivo* transcriptomic data (14) (**Fig. 7A**). A triple mutant targeting the uninduced transporters *bglF*, *chbB*, and *agaV*, with predicted substrates β-glucoside, cellobiose, and N-acetyl-galactosamine, did not exhibit any fitness defect in co-challenge experiments (**Fig. S7**), suggesting these importers are dispensable in the urinary tract.

**Fig. 7.**
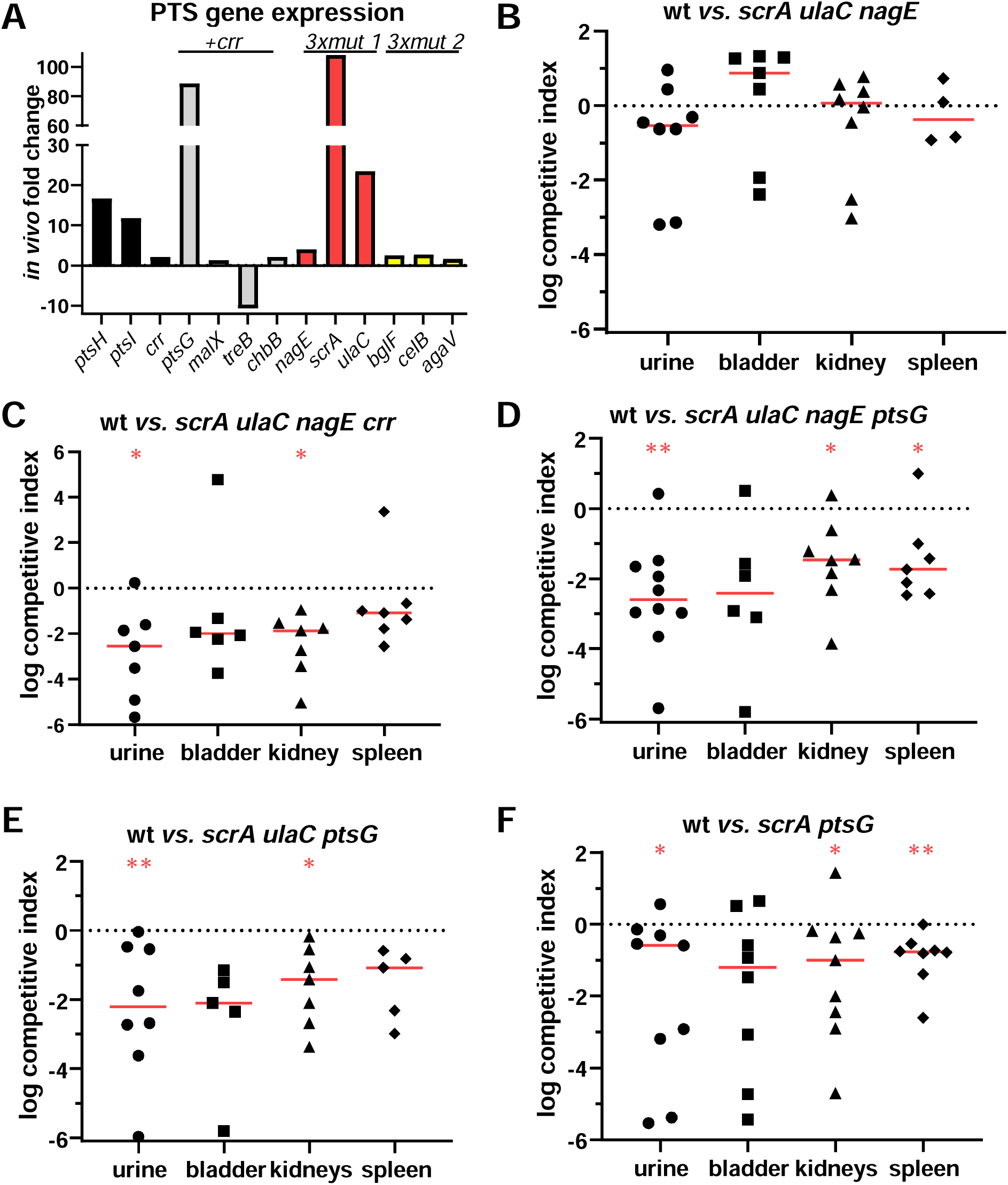
PTS multi-mutant co-challenges. **A,** *In vivo* differential expression was used to prioritize mutant construction. Red and yellow bars correlate with the arrows in Fig. 5. **B,** a combinatorial triple mutant constructed based on higher induction during experimental UTI (red bars) did not have a competitive disadvantage. **C,** quadruple mutant with *crr* recapitulated *ptsH* phenotype. **D,** quadruple mutant with *ptsG* also was similar to *ptsH* phenotype. **E,** removing *nagE* preserved the *ptsH* phenotype. **F,** an *scrA ptsG* double mutant had a much weaker defect, emphasizing the combinatorial contribution of different PTS transporters to fitness during experimental UTI. Horizontal lines show medians. Dashed line indicates equal fitness of wild type and mutant (log CI= 0). \**P* < 0.05; \*\**P* < 0.01; one sample Wilcoxon test *vs.* a theoretical median of 0.

We next tested a triple mutant targeting the highly induced genes *nagE*, *scrA*, and *ulaC*, which encode predicted transporters for N-acetylglucosamine, sucrose, and L-ascorbate, respectively. This strain also showed no significant *in vivo* defect (**Fig. 7B**). However, the addition of a fourth mutation in *crr* resulted in a composite fitness defect, with significantly reduced recovery from urine and kidneys (median log_10_ CI −2.55, and −1.87, respectively) (**Fig. 7C**). Of the importers predicted to work with Crr, glucose-specific EIIBC component *ptsG* transcripts were the only ones induced during experimental UTI in mice (**Fig. 7A**). Replacing *crr* with a *ptsG* mutation produced a similar result, with significant attenuation in urine, kidneys, and spleen (median log_10_ CI −2.60, −1.47, and −1.74, respectively) (**Fig. 7D**). Both quadruple mutants phenocopied the fitness loss seen for the *ptsH* mutant (**Fig. 2C**), indicating that the substrates imported by these systems collectively contribute to *in vivo* fitness. CFU recovery for all PTS mutant co-challenges are shown in **Fig. S8**.

To determine the minimal functional set of PTS importers needed for urinary tract fitness, we progressively removed individual genes from the *nagE*/*scrA*/*ulaC*/*ptsG* quadruple mutant. After restoring wild type *nagE*, the least induced gene (**Fig. 7A**), the defective phenotype of triple mutant *scrA*/*ulaC*/*ptsG* was not altered (significant median log_10_ CI of −2.22 and −1.42 in the urine and kidneys, respectively) (**Fig. 7E**). However, further removal of the *ulaC* mutation reduced the magnitude of the competitive defect, although the mutant remained significantly attenuated in urine, kidneys, and spleen (median log_10_ CI −0.59, −1.00, and −0.77, respectively) (**Fig. 7F**). These results suggest that import mediated by *ulaC*, *scrA*, and *ptsG*, collectively, are the cause of the PTS-dependent *in vivo* defect first observed with the *ptsH* and *ptsI* mutants (**Fig. 2 C-D**).

Although we identified *ulaC*, *scrA*, and *ptsG* as important contributors to *in vivo* fitness, only PtsG had an experimentally confirmed substrate in *P. mirabilis* (glucose, **Fig. 1B**). UlaC and ScrA are predicted to import L-ascorbate and sucrose, respectively. However, wild-type HI4320 failed to utilize 10 mM L-ascorbate as a carbon source under aerobic conditions (**Fig. S9A**) and instead showed steadily declining optical density compared to control media lacking a carbon source. *E. coli* has been reported to ferment L-ascorbate under anaerobic conditions (53). However, an anaerobic atmosphere did not allow *P. mirabilis* growth on ascorbate (**Fig. S9B**), further suggesting that this substrate is not imported by HI4320. While the PMI1775-1777 operon appears to encode a PTS IIA, IIB, IIC importer, closer examination of this locus in other databases did not yield viable alternative substrates. TransportDB listed fructose as the substrate, but HI4320 did not grow on D-fructose in the Biolog PM1 carbon source panel (**Table S2**).

Likewise, we found that sucrose was not a functional carbon source for strain HI4320 (**Fig. 6E**), raising questions about the annotated role of ScrA. The gene encoding *scrA*, PMI3515, appears to be part of a four-gene operon comprising PMI3514-17 (**Fig. S9C**). The protein encoded by PMI3514 has 47% identity and 67% similarity to *E. coli* MG1655 MurQ, which is an *N*-acetylmuramic acid (MurNAc) 6-phosphate etherase. In *E. coli*, the next gene encodes MurP, a PTS enzyme IICB component that imports MurNAc and shares 53% similarity to HI4320 ScrA. PMI3516 encodes a protein that looks like transcriptional regulator MurR, although in *E. coli*, *murR* is divergently transcribed from the *mur* operon. However, the last genes in each operon (PMI3517 or *yfeW*) are dissimilar. We therefore hypothesized that ScrA imports MurNAc. Despite the similarities, HI4320 was unable to grow using 0.2% MurNAc as a sole carbon source (**Fig. S9D**). Thus, although both *ulaC* and *scrA* contributed to *P. mirabilis* fitness and are induced during experimental UTI, the substrates for both remain to be determined. These results highlight the need for experimental validation of sugar transporter function in *P. mirabilis*, as many annotations based on *E. coli* homology may not accurately reflect substrate specificity in this organism.

### Glucosuria enhances *P. mirabilis* colonization and increases the severity of infection

The *in vivo* defects observed with *ptsH* and *ptsI* mutants reinforce prior studies demonstrating that carbohydrate metabolism is critical for *P. mirabilis* fitness during urinary tract infection (14, 15). Although sugars are not typically considered a major carbon source in urine, they are clearly accessible to *P. mirabilis* during experimental UTI. We hypothesized that increasing urinary sugar levels would intensify the fitness disadvantage observed for the *ptsH* mutant. We accomplished this using the sodium-glucose cotransporter 2 (SGLT2) inhibitor dapagliflozin, which reduces renal glucose reabsorption leading to increased glucose concentrations in urine (54).

As expected, administration of dapagliflozin via drinking water increased glucosuria by more than 20-fold within 24 h in female CBA/J mice, raising the mean glucose from 375 to 14,288 mg/dL (**Fig. 8A**). Elevated glucosuria remained for the duration of the 7-day experiment, falling slightly to 8,311 mg/dL by day 7. Following infection with wild-type HI4320, urinary glucose levels remained relatively stable in mice that received normal drinking water over 7 days (**Fig. 8B**). Interestingly, over time, two of the mice inoculated with *P. mirabilis* showed glucose levels that declined by more than half over the course of the experiment; one animal strikingly had an over 90% reduction from baseline by day 3 (77.5 *vs.* 7.21 mg/dL) that further fell below the limit of detection by day 7. Dapagliflozin-treated mice showed increased urinary glucose similar to the pilot study, marked by a stark reduction in urinary glucose on day 3 post-inoculation, suggestive of bacterial glucose consumption (**Fig. 8B**). Additionally, 20% (n=2) of dapagliflozin-treated animals reached humane endpoints before or at the scheduled termination of the study. This was consistent with higher median bacterial burdens in the urine, particularly on day 3 post-inoculation (**Fig. 8C**). At 7 days post-inoculation, kidney colonization was significantly increased in the dapagliflozin group (**Fig. 8D**); bladder colonization was also trending toward a significant increase (*P*=0.0635, n=4 in the treated group). These results demonstrate that SGLT2 inhibition effectively induces glucosuria in CBA/J mice and promotes enhanced colonization and disease severity during *P. mirabilis* UTI.

**Fig. 8.**
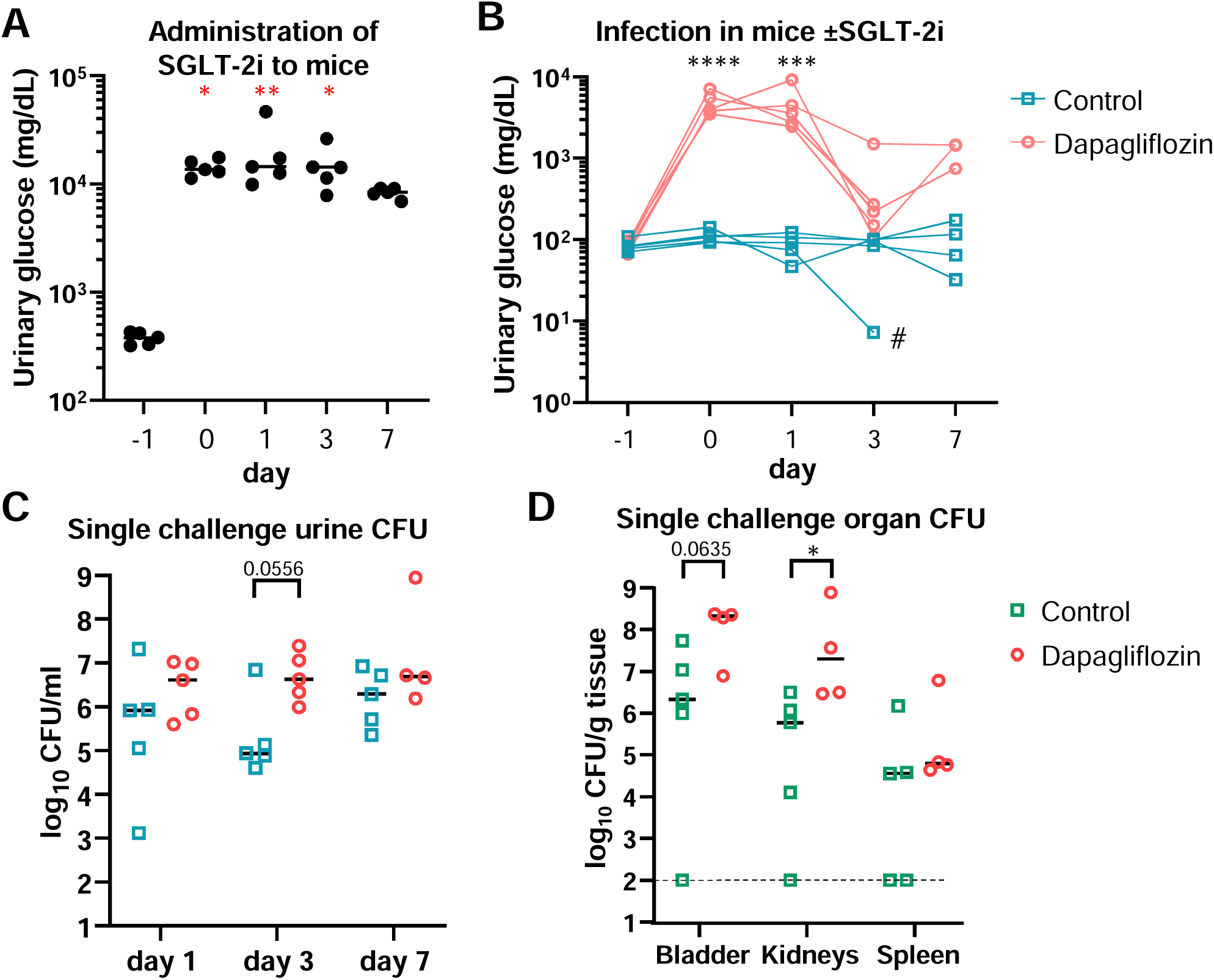
Experimental *P. mirabilis* UTI during SGLT-2i treatment. **A,** Measurement of urinary glucose. Administration of SGLT-2 inhibitor dapagliflozin to female CBA/J mice via drinking water on “day −1” resulted in increased glucose excretion via urine. \**P*<0.05, \*\**P*<0.01 *vs.* day −1, two-way ANOVA with Dunnett’s multiple comparisons test. **B-D,** Experimental inoculation of *P. mirabilis* HI4320 on day zero in mice with or without dapagliflozin treatment (n = 5/group). Of the five mice in the dapagliflozin group, one mouse died at day 6 and one was moribund at day 7 (unable to collect enough urine for glucose measurement). **B,** Urinary glucose. Lines connect data from individual mice. Glucose levels fell somewhat in infected mice, and the effect was more apparent in the dapagliflozin-treated mice. # indicates one control mouse with undetectable glucose at day 7. \*\*\**P*<0.001, \*\*\*\**P*<0.0001 for dapagliflozin-treated mice *vs.* day −1, mixed-effects analysis with Dunnett’s multiple comparisons test. **C,** Bacterial burden in urine. Median CFU recovery was higher in dapagliflozin-treated mice, although with 5 mice the difference was not statistically significant. **D**, Bacterial burden in organs at experimental endpoint (7 dpi). Dashed line indicates limit of detection. C and D, \**P*<0.05; for 0.05<*P*<0.1, exact value shown; Mann-Whitney test.

Due to the increased morbidity observed during infection of dapagliflozin-treated mice, we next tested the fitness of the *ptsH* mutant in a 3-day co-challenge experiment, timed to coincide with peak glucose utilization and elevated urinary colonization (**Fig. 8B-C**). As expected, urinary glucose levels remained elevated in dapagliflozin-treated mice and declined by day 3, again suggesting bacterial consumption (**Fig. 9A**). Consistent with previous findings, dapagliflozin-treated mice exhibited higher overall urinary colonization (**Fig. S10A**), and the fitness defect of the *ptsH* mutant was significant on day 2 in treated mice but not in controls (median log_10_ CI −0.64 *vs*. −0.38 control) (**Fig. 9B**). By day 3, the *ptsH* mutant showed a significant fitness defect in the urine of both groups (median log_10_ CI −1.54 *vs*. −0.80 control), and this was also observed in the bladder (−1.59 *vs.* −0.83 control) (**Fig. 9C**). Interestingly, although the fitness defect of the *ptsH* mutant in dapagliflozin-treated mice was not significantly different from control mice in the kidneys (median log_10_ CI −1.04 vs. −1.71 control) and spleens (median log_10_ CI −0.99 vs. −1.22 control), in both organs, the CI reached statistical significance only for dapagliflozin-treated mice. This is because bacteria were recovered from all mice in the dapagliflozin group, whereas many mice in the control group did not have measurable ascending (n = 5; 50%) or disseminated (n = 7; 70%) infection (**Fig. S10A-B**). Adding up the total CFU (wild type plus *ptsH*) recovered from each mouse showed that overall bacterial burden was significantly higher in mice treated with dapagliflozin (**Fig. S10C**). When glucose levels were correlated with bacterial burdens on day 3, a strong inverse relationship was observed in control mice across urine, bladder, and kidney samples (R² = 0.28, 0.64, and 0.84, respectively) (**Fig. 10A, C, E**). No such trend was seen in the dapagliflozin-treated group (**Fig 10 B, D, F**), suggesting that hyperglucosuria alters the physiological responses of *P. mirabilis* during experimental UTI. These findings support the use of dapagliflozin to model glucosuria in CBA/J mice and demonstrate that elevated urinary glucose exacerbates *P. mirabilis* colonization and infection severity.

**Fig. 9.**
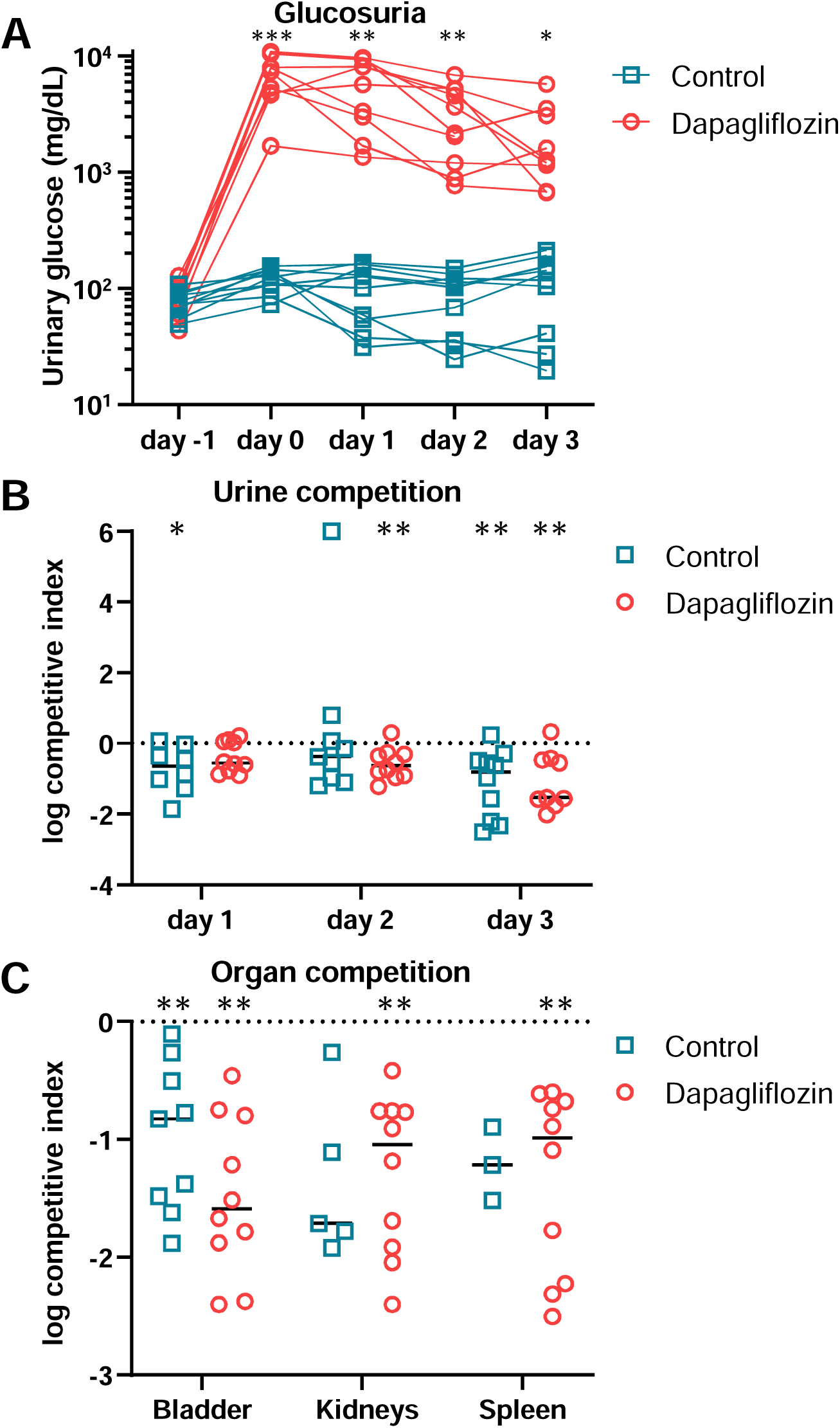
Wild type *vs*. *ptsH* co-challenge during SGLT-2i treatment. Twenty mice, half receiving dapagliflozin and half receiving normal water, were administered a 1:1 mixture of wild type and *ptsH* mutant bacteria. A, urinary glucose. Lines connect urinary glucose levels over time in individual mice. \**P* <0.05, \*\**P* < 0.01, \*\*\**P*<0.001 control *vs.* dapagliflozin, mixed-effects analysis with Šidák’s multiple comparisons test. B-C, competitive indices. Horizontal lines show medians. \**P* <0.05, \*\**P* < 0.01 *vs.* theoretical median of log CI = 0, one sample Wilcoxon test. Comparisons between control and dapagliflozin groups were not significant (Multiple Mann-Whitney tests with Holm-Šídák correction). B, urinary competitive index measured over time. C, organs at experimental endpoint (3 dpi).

**Fig. 10.**
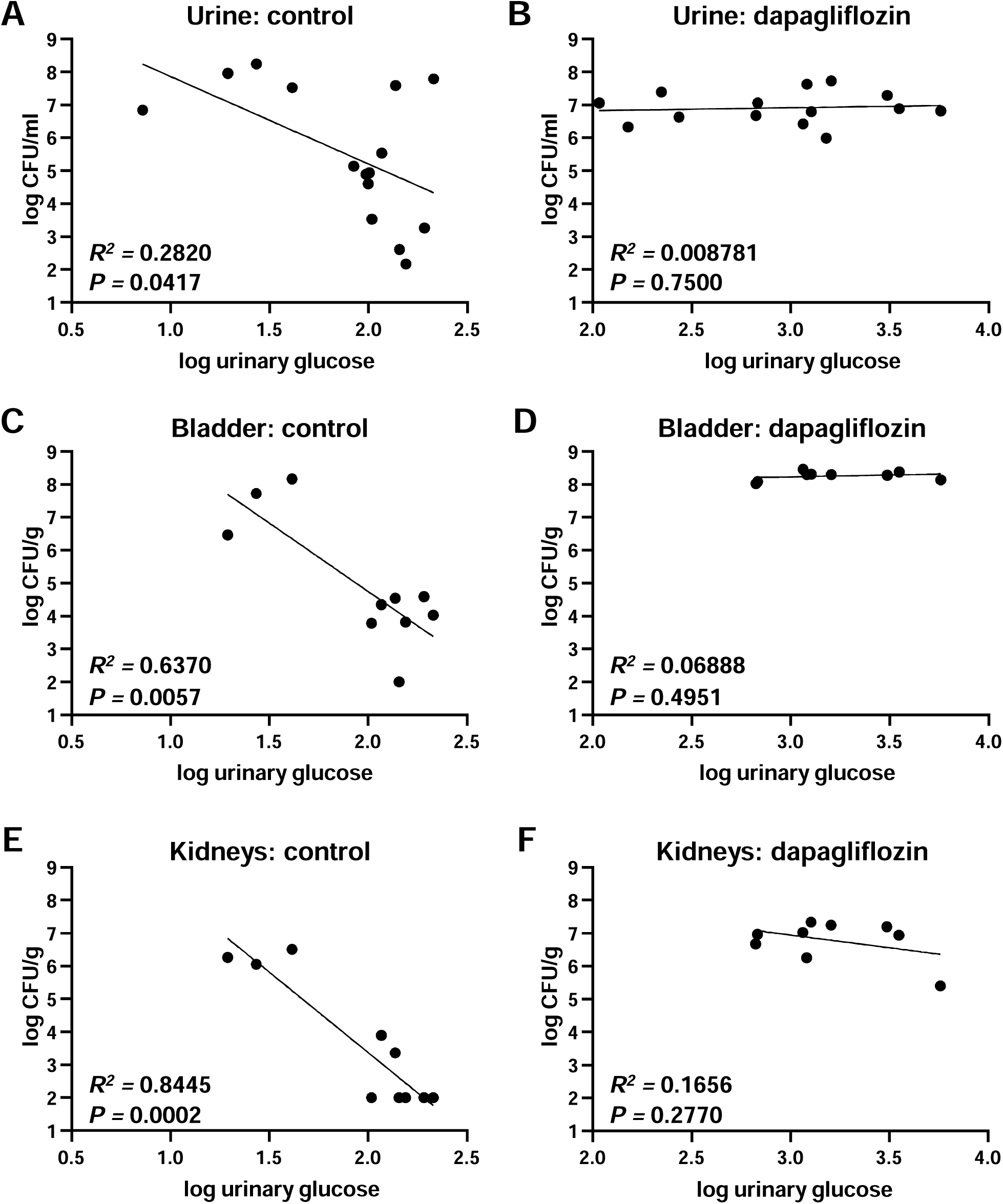
Inverse correlation between glucosuria and CFU recovery. (**A, C, E**) but not during SGLT-2i-induced hyperglucosuria (**B, D, F**) at 3 dpi. Data were compiled from wild type single challenge and wild type *vs. ptsH* co-challenge (total CFU; limit of detection = 100 for organs). Lines indicate simple linear regression.

## DISCUSSION

The metabolic strategies employed by bacterial pathogens to colonize the urinary tract differ by species, reflecting adaptations to specific host niches. UPEC relies primarily on amino acid metabolism *in vivo*, with carbohydrate utilization playing a minor role (11, 15, 18). In contrast, prior transcriptomic studies have shown that *P. mirabilis* upregulates glycolytic enzymes and sugar transporters during UTI, in addition to amino acids, suggesting a greater reliance on sugar-derived carbon sources (14). Here, we directly assessed the importance of sugar import systems in *P. mirabilis* by generating 47 targeted transporter mutants and evaluating both their *in vitro* growth and *in vivo* fitness. Several mutants displayed *in vivo* fitness defects despite lacking observable phenotypes in standard growth conditions, indicating that specific import systems contribute to pathogenesis in a context-dependent manner.

*In vivo* pooled mutant screens are inherently limited by physiological bottlenecks in the ascending UTI model, including for *P. mirabilis* (42). Although our In-seq experiments were designed to remain within established bottleneck constraints, we nonetheless observed clear founder effects in a subset of mice, where 1-3 mutants disproportionately dominated sequencing reads from individual organs. Similar stochastic population dynamics have been reported in UPEC infection models, even under carefully controlled conditions (55–57). Despite these limitations, all fitness determinants identified by In-seq were confirmed in a traditional 7-day co-challenge model, exceeding validation rates reported in comparable *E. coli* studies (56). These findings underscore both the challenges of pooled *in vivo* approaches and the robustness of our experimental design and follow-up strategy.

The Major Facilitator Superfamily (MFS) comprises a large and diverse group of membrane transport proteins characterized by 12 transmembrane helices and driven by electrochemical gradients. MFS transporters are present in both prokaryotes and eukaryotes; for example, glucose uptake in humans is mediated by members of this family (58). In this study, *xapB*, an MFS transporter, was identified as important for *P. mirabilis* fitness during infection. Although XapB is annotated as a xanthosine permease based on homology to *E. coli* (44), its substrate specificity in *P. mirabilis* remains unverified. Prior studies support a role for nucleoside metabolism in *P. mirabilis* polymicrobial pathogenesis; loss of xanthine-guanine phosphoribosyltransferase (*gpt*) and purine-nucleoside phosphorylase (*deoD*) impaired *in vivo* fitness during co-infection with *Providencia stuartii* (16). However, we found that strain HI4320 could not use xanthosine as a sole carbon or nitrogen source under the conditions we tested, and guanosine uptake occurred independently of XapB, validated in a *guaA/xapB* double mutant. *P. mirabilis* HI4320 encodes nine additional genes that are annotated as nucleoside importers, suggesting there are multiple routes of entry for these molecules.

Importantly, the *xapB* mutant displayed a significant defect during experimental UTI, suggesting that the imported substrate(s) is likely non-redundant. In this study, we did not initially set out to study nucleoside import; *xapB* was included in our 47 mutant panel because it was annotated as a generic sugar importer by TransportDB at the start of the project. Interestingly, over the course of this work, TransportDB updated the annotation for *xapB* at least twice, first to generic nucleoside transport, and then to melibiose importer *melB.* Melibiose is a plant disaccharide that did not support growth of HI4320 in our carbon source testing, leading us to conclude that this revised annotation is also likely incorrect. These findings highlight a recurring challenge in microbiological research: reliance on gene annotations inferred from *E. coli* often fail to recapitulate actual function.

Similar annotation discrepancies were observed for multiple phosphotransferase system (PTS) family transporters in our study. PtsH and PtsI are conserved upstream components of the PTS, a multi-component phosphorelay essential for sugar uptake and carbon catabolite repression (24). Both *ptsH* and *ptsI* mutants exhibited reproducible defects in a murine model of ascending UTI, consistent with a central role for glycolysis in *P. mirabilis* pathogenesis and contrasting with UPEC and other uropathogens (14, 15, 59). Even so, because PtsH and PtsI interact with many sugar-specific components, these findings did not identify the specific substrates responsible for fitness. Here, we found that the three most-induced substrate-specific PTS genes during experimental *P. mirabilis* UTI in mice, *ptsG*, *scrA*, and *ulaC*, combined to recapitulate the *ptsH* mutant phenotype. However, *in vitro* substrate validation for two of these transporters was inconsistent with predicted annotations. HI4320 was unable to use either sucrose, predicted to be imported by ScrA, or L-ascorbate, predicted to be imported by UlaC, as a carbon source. Nor did ScrA support growth on MurNAc in follow-up experiments despite homology with the *E. coli mur* locus. Likewise, HI4320 failed to grow on predicted PTS substrates maltose, cellobiose, and β-glucosides. Incidentally, although *E. coli* was initially shown to utilize cellobiose, later work indicated the more physiologically likely substrate is chitobiose (60). Because the *chbB* mutant was not found to be important for *P. mirabilis* fitness during experimental UTI (**Fig. S7**), we did not investigate chitobiose further. Overall, only 4/9 of the KEGG-predicted PTS substrates for *P. mirabilis* verified experimentally. Notably, our findings are consistent with Biolog data for *P. mirabilis* isolates obtained from broiler chickens (61).

Beyond sugar transport, the PtsHI phosphorelay appears to influence nitrogen metabolism. The *ptsH* mutant showed defects when grown on ammonia, urea, and several amino acids, suggesting cross-regulation between carbon and nitrogen pathways. Likewise, both *ptsH* and *ptsI* mutants displayed a more pronounced defect on glycerol compared with the more substrate-specific *crr* and *ptsG* mutants. In *E. coli*, the PTS^Ntr^ system coordinates nitrogen assimilation by sensing glutamine and α-ketoglutarate (28, 62), but has not been characterized in *P. mirabilis*. Our findings raise the likelihood of similar regulatory integration.

While the urinary tract is generally sugar-poor, elevated glucose levels occur in specific conditions such as diabetes or treatment with SGLT2 inhibitors. Diabetic individuals have increased UTI susceptibility, and glucosuria is common in both humans and mouse models (59, 63). SGLT2 inhibitors, such as dapagliflozin, reduce renal glucose reabsorption, leading to elevated urinary glucose. Although debated in clinical studies (64, 65), preclinical studies consistently show worsened UTI outcomes under glucosuric conditions, including increased bacterial burden and dissemination (22, 66). We observed similar results for *P. mirabilis* infection, where dapagliflozin treatment led to elevated colonization and exacerbated disease. Despite these findings, the *ptsH* mutant did not demonstrate an enhanced competitive defect in urine under glucosuric conditions, as predicted. One explanation for this result is that additional sugar import systems contribute to fitness during UTI, thereby partially masking the effect of *ptsH* loss under glucosuric conditions. Consistent with this idea, growth of the *ptsH* mutant in minimal medium with glucose was slowed but not abolished, indicating that glucose can still enter the cell through alternative pathways. Several transporters, including *crr*, exhibited borderline effects on *in vivo* fitness in the In-seq screen, suggesting that their contributions may become apparent only in specific metabolic contexts. It is also possible that loss of *ptsH* induces compensatory metabolic or regulatory responses that mitigate fitness defects when glucose availability is increased.

The connection between glucosuria and UTI severity likely involves more than nutrient availability. Hyperglycemia impairs innate immunity, including reduced neutrophil responses and cytokine signaling in the kidney (22, 67). We observed an inverse correlation between urinary glucose and bacterial burden in untreated mice, but this trend was lost in dapagliflozin-treated animals, likely reflecting both increased glucose consumption and altered host-pathogen interactions. Mouse background also matters; others have reported that C57BL/6J mice treated with dapagliflozin had similar bladder and urine colonization by UPEC (23). C3H/HeOuJ mice had to be administered 10 mg/kg of dapagliflozin to see increased UPEC burden in bladder and kidneys, and results were dependent on bacterial strain (23). Mice genetically engineered to develop type 2 diabetes (*db*/*db*) also have increased UTI risk that is correlated with altered innate immune markers (68). However, dapagliflozin-treated CBA/J mice were shown to have increased UPEC and *K. pneumoniae* colonization at chronic and acute timepoints (22). Interestingly, while UPEC infections in this model could be sustained for 7 days, *P. mirabilis* infection resulted in increased morbidity at this time point, consistent with this species’ increased contribution to UTI in complicated backgrounds.

In conclusion, our findings provide mechanistic insight into the metabolic requirements of *P. mirabilis* during UTI. Sugar import systems, particularly PTS transporters, are essential for *in vivo* fitness, and substrate annotations based on *E. coli* must be experimentally validated. Our data reveal regulatory crosstalk between carbon and nitrogen metabolism and demonstrate that host metabolic conditions, such as glucosuria, exacerbate infection severity. These results have implications for managing UTI in individuals with metabolic disease and highlight the need for continued investigation into the roles of uncharacterized transporters and their regulation in *P. mirabilis* pathogenesis. They also underscore a broader gap in our understanding of the *P. mirabilis* genome and the limitations of relying on comparative functional annotation. Future work will focus on identifying the specific substrates imported by fitness-contributing transporters, confirming the presence of these metabolites in the urinary tract, and determining how *P. mirabilis* accesses these nutrient sources during infection.

## METHODS

### Bacterial strains and culture conditions

*P. mirabilis* strain HI4320 was isolated from the urine of an elderly female nursing home patient with a long-term (≥30 days) indwelling catheter (3, 37, 69). *E. coli* TOP10 (Thermo Fisher) was used for plasmid construction and maintenance. Bacteria were routinely cultured at 37°C in lysogeny broth (LB; per liter: 10 g tryptone, 5 g yeast extract, 0.5 g NaCl) with aeration or on LB solidified with 1.5% agar. As needed, antibiotic selection was used as follows (µg/mL): kanamycin 25, ampicillin 50, chloramphenicol 20. For experiments requiring minimal, chemically defined media, Minimal A was used (70). The carbon source in Minimal A was 0.2% glycerol unless otherwise specified. All strains used in this study are listed in **Table S3.**

### Mutant construction

All mutants were constructed using a *P. mirabilis*-tailored version of targetron insertional mutagenesis (39, 71). Briefly, stable chromosomal mutations were constructed using a synthesized 353 bp group II intron fragment (eBlocks, Integrated DNA Technologies) that specifically targeted each gene designed using the ClosTron prediction algorithm (72). Reprogrammed intron fragments were cloned into pACD4K-CloxP using NEBuilder HiFi DNA Assembly master mix (New England Biolabs) with primers designed to amplify vector or intron templates and confirmed by DNA sequencing (Eurofins). Targetron-containing plasmids and a source for T7 polymerase, pAR1219 (73), were introduced into *P. mirabilis* HI4320 using electroporation and induced to jump into the specified genes. Insertional mutations in kanamycin-resistant mutants were confirmed using PCR. To construct multiple mutations in the same background, the kanamycin resistance gene in the initial insertion was removed using *cre*/lox recombination to create a markerless mutant (39, 74). Targetron sequences are listed in **Table S4.**

### Murine model of ascending UTI

Bacterial fitness during UTI was assessed using a well-established mouse model (45, 75, 76). Briefly, overnight cultures of *P. mirabilis* were diluted in LB to OD_600_ = 0.092-0.094 (∼2 × 10^8^ CFU/mL). For co-challenge experiments, wild type and mutant bacteria were mixed 1:1. Ten female CBA/J mice, aged 5-6 weeks (Jackson Laboratory), were transurethrally inoculated with 50 µL of this 1:1 mixture (10^7^ CFU/mouse) over 30 s using a Harvard pump. At 7 days post-inoculation, urine was collected; mice were euthanized; and bladders, kidneys, and spleens were harvested. Organs were homogenized and plated to quantify CFU; mutants were distinguished from wild-type colonies using kanamycin. Competitive indices were calculated for each site by comparing the ratio of output wild type and mutant to the ratio of input bacteria (45, 77). For sites with no recovered CFU, the limit of detection for urine was set to 20 and for organs 100. Statistical significance of competitive indices was calculated using the Wilcoxon signed rank test.

### Pooled transporter mutant murine challenge

To measure the relative contributions of sugar transporters to UTI, 47 targetron mutants were individually cultured and the density adjusted as described above. 23 (ABC + MFS, group 1) or 24 (PTS + Others, group 2) strains were mixed together in equal volume, and 15 mice/group were transurethrally inoculated with 50 µL (∼10^7^) CFU of either mixture. A 1 mL aliquot of each input was directly pelleted for DNA purification and a second aliquot was plated to quantify CFU (group 1, 2.51 × 10^8^ CFU/mL; group 2, 2.68 × 10^8^ CFU/mL). After 24 h, urine was collected and diluted to 250 µL, mice were euthanized, and bladders and kidneys were collected. Organs were homogenized (Omni International) in 2 mL phosphate-buffered saline (PBS); a portion of organs and urine was dilution-plated to determine output CFU, and the remainder was spread-plated on LB agar containing kanamycin. The next day, colonies from each plate were swabbed into 10 mL of PBS, pelleted, and frozen. Colonies from the plated input samples were also collected as a control for growth on agar (input spiral, insp). Chromosomal DNA was purified from inputs and urine, bladder, and kidney outputs using the Wizard Genomic DNA Purification Kit (Promega) and quantified using a Qubit fluorometer (Invitrogen).

### Insertional site sequencing (In-seq)

Targetron junctions were enriched and prepared for high-throughput sequencing with modification of established protocols (78, 79). Briefly, we used the NEBNext Ultra II FS DNA Library Prep with Sample Purification Beads kit to fragment DNA into 200-450 bp lengths and ligate sequencing adaptors with sequences modified to be in line with lower %GC content in *P. mirabilis* (TA_Adaptor_Top and TA_Adaptor_Bottom). Targetron-gene junctions were enriched by PCR using primers Targetron_enrich_For and Transposon_enrich_Rev (**Table S5**). Samples (n = 78) were barcoded by indexing PCR to label each specific library using NEBNext Multiplex Oligos for Illumina (Dual Index Primers Set 1) (**Table S1**). PCR products were measured by Qubit and submitted to the University of Michigan Advanced Genomics Core for Illumina sequencing (150 nt, paired-end).

### In-seq analysis

Mutant fitness in mice was assessed by quantifying the proportion of each of the 47 targetron-gene junctions in the input and output sequences. Identification and quantification of targetron junctions was conducted by the University of Michigan Medical School’s Bioinformatics Core. Filtering, trimming, and deduplication of reads was accomplished using an established Tn-seq pipeline (16, 80), and subsequent steps were adapted for the relatively small number of targetron insertions. BLASTN with the 3′ end of the targetron as query was used to identify targetron-containing sequences consisting of at least 100 nt at 100% identity on the plus strand. The 19 nt immediately following the targetron sequence were used to uniquely identify the locus of gene insertion for each targetron (*i.e.*, 47 input mutants). A counts matrix was generated from the 1,739,134 unique 19 nt reads. Features which were members of each individual group (1 or 2) were input into EdgeR, a limma-based R package which is able to deal with group sizes of only one sample in a differential enrichment calculation (81). Deduplicated library sizes were used for depth normalization.

### Diabetic UTI murine model

To assess *P. mirabilis* UTI during glucosuria, we adapted the method from Nishitani *et al.* (82). The SGLT-2 inhibitor dapagliflozin (MedChem Express) was dissolved in ethanol (125 mg/ml), then diluted in Ann Arbor city water to a final concentration of 0.02 mg/ml. Female CBA/J mice were administered dapagliflozin via drinking water *ad libitum* beginning 24 h before bacterial inoculation. Bacteria were prepared as described above. Urine was collected at specified intervals and glucose was quantified using an Infinity glucose hexokinase assay (Thermo Fisher) according to the manufacturer’s instructions. Aliquots of urine from the same time points were diluted and plated to quantify CFU, and bacterial burden in organs was assessed as described above.

### Growth curves

Bacterial growth over time (24 h) was measured in triplicate by recording the optical density at 600 nm (OD_600_) at 15 min intervals using a Bioscreen C set to 37°C with continuous shaking. Chemical complementation of growth defects was performed in Minimal A medium with 0.2% glycerol as the carbon source unless otherwise specified. Anaerobic growth curves were conducted in an anaerobic chamber (Coy Lab Products, Grass Lake, MI) at 37°C under an atmosphere of 5% H_2_, 5% CO_2_, and 90% N_2_. OD_600_ of 96-well plates was measured every 10 mins for 48 h in a microplate stacking device (BioStack 2WR) with coupled absorbance reader (Powerwave HT, BioTek Instruments). Doubling times for the 47 transporter mutants were calculated from three biological replicates using the AMiGA software package in [R] (83) from 0.5 to 12 hours. Doubling time was computed as ln(2) multiplied by the inverse of the maximum specific growth rate. To identify potential sugar transporter substrates, Biolog Phenotype MicroArray plates PM1-2 (carbon sources) and PM3 (nitrogen sources) were inoculated with wild-type or mutant *P. mirabilis* and assayed for growth every 10 min for 24 h using a LogPhase 600 Microbiology Reader (BioTek) with shaking. Biolog cultures were conducted using the manufacturer’s recommended medium with the respiration substrate omitted.

### Swarming motility

Swarming motility experiments were conducted as previously described (84). Briefly, 5 µl of a logarithmic-phase culture was added to the center of an LB (10 g/L NaCl) agar plate, allowed to dry, and incubated at 30°C for 16 h, after which the swarming radius was measured.

### Statistical analysis

All graphs were plotted and statistics calculated using GraphPad Prism 10. Statistical tests and significance values used for each experiment are indicated in figure legends. Error bars show SD unless otherwise indicated.

### Ethics approval

Animal experiments were approved by the University of Michigan Medical School Institutional Animal Care and Use Committee, protocol number PRO00010856. During catheterization procedures, mice were anesthetized by intraperitoneal injection of ketamine/xylazine. Mice were euthanized by inhalant isoflurane anesthetic overdose prior to organ removal.

## Data availability

Sequencing and analysis data from the pooled transporter In-seq experiments are available at GEO accession #GSE244606. All strains and plasmids are available upon reasonable request.

## Acknowledgements

We thank Valerie Forsyth for assistance with primer design for insertion sequencing, coordination with core facilities, and adapting legacy protocols for updated kits. Santosh Paudel provided support with urine collections and animal infections. Devra Deleston assisted with some growth curve experiments. We are grateful to Margaret Stiner and the ULAM Technical Services team for their support with murine co-challenge infections and the diabetic model. We thank Martin Myers from the Michigan Diabetes Research Center for advice in modeling glucosuria. We also acknowledge support from Rebecca Tagett and Weisheng Wu at the Bioinformatics Core of the University of Michigan Medical School’s Biomedical Research Core Facilities (RRID:SCR_019168).

## SUPPORTING INFORMATION

### Supplemental Figure Legends

**Supplemental Fig. S1.**
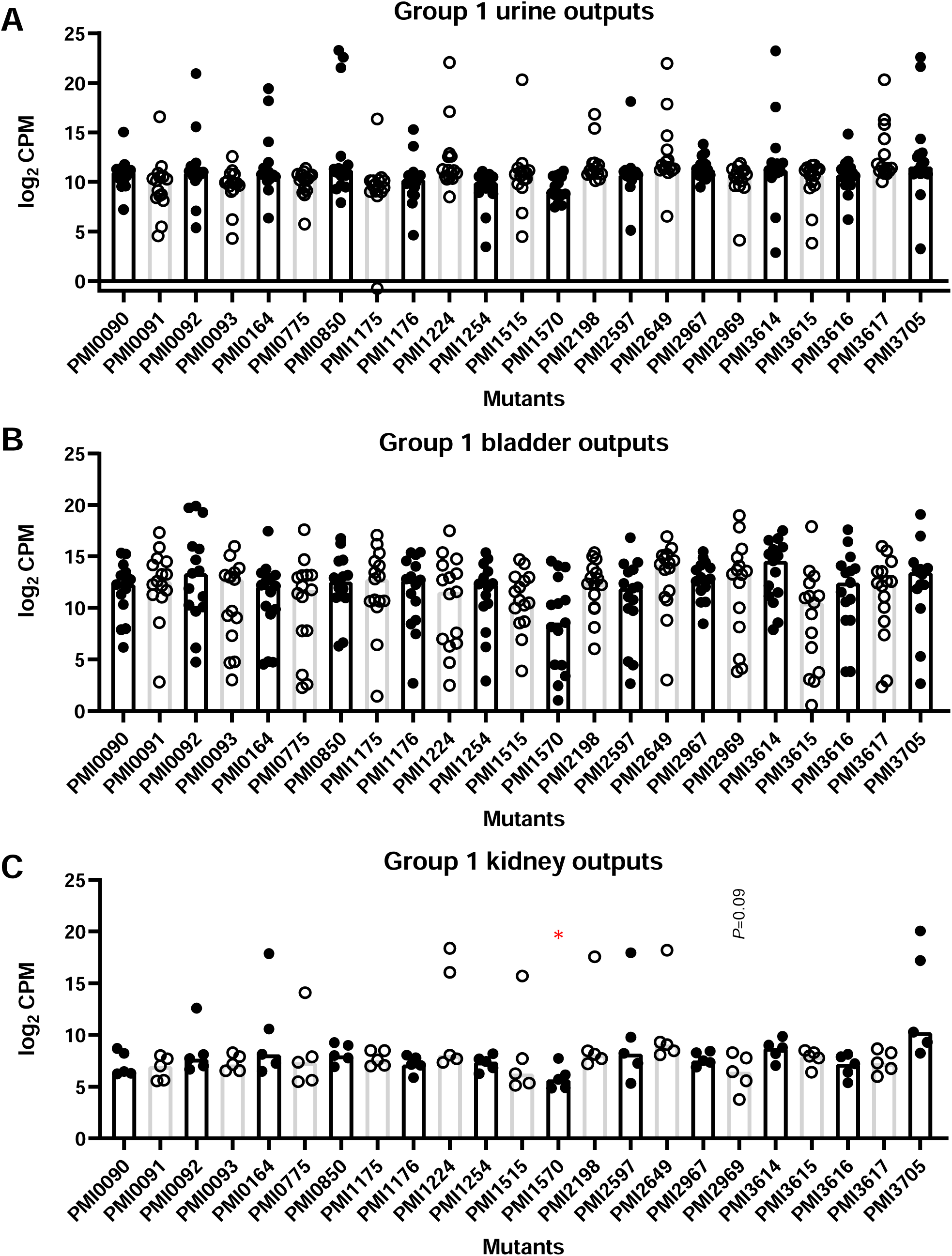
Group 1 (ABC and MFS) mutant recovery from mice, 24 h post-infection. **A**, urine; **B**, bladder; **C**, kidneys. Each data point represents sequencing reads obtained from one mouse. \**P* < 0.05. Exact *P* values shown when 0.1 > *P* > 0.05.

**Supplemental Fig. S2.**
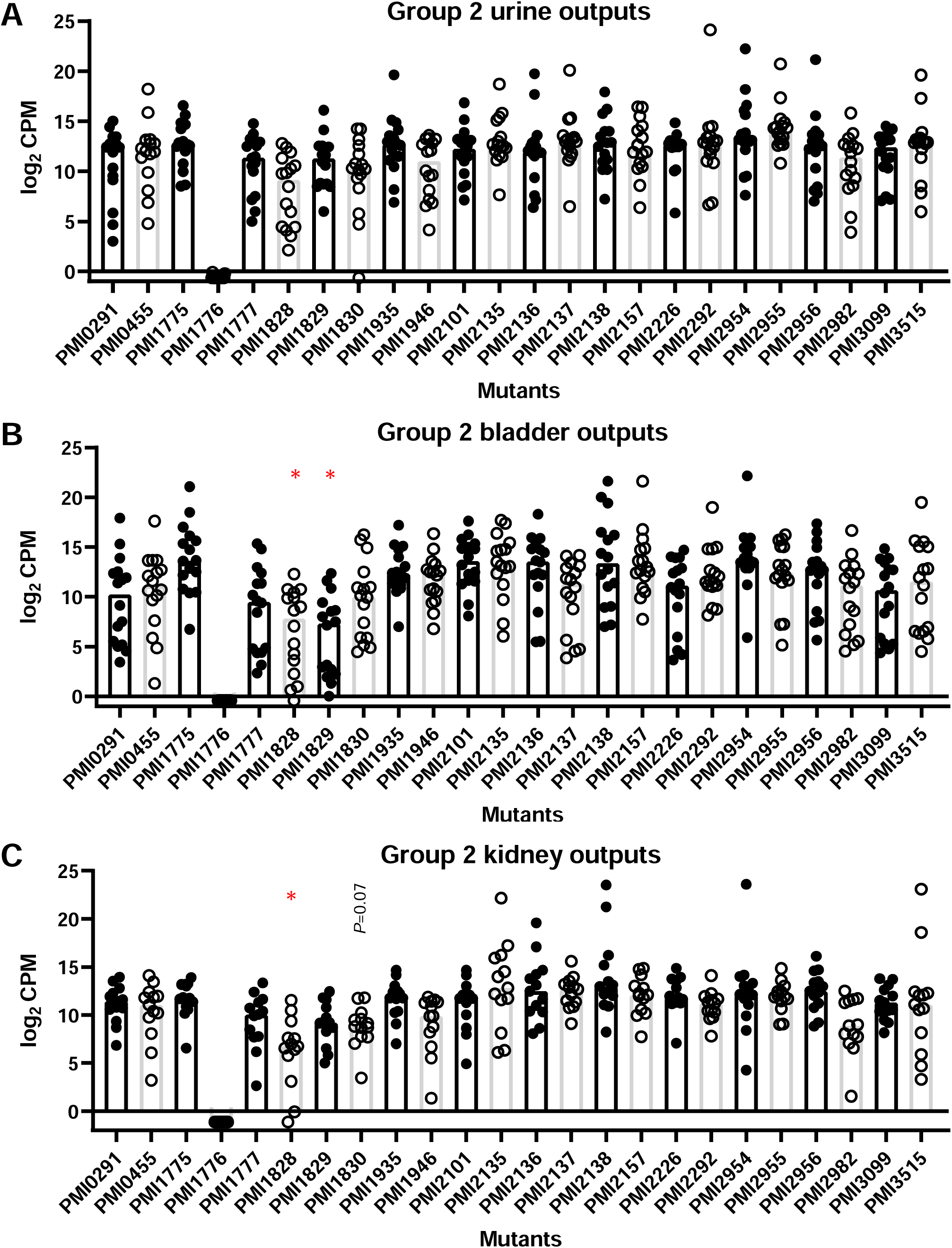
Group 2 (PTS and Others) mutant recovery from mice, 24 h post-infection. **A**, urine; **B**, bladder; **C**, kidneys. PMI1776 is missing because PMI1176 was inadvertently inoculated instead. Each data point represents sequencing reads obtained from one mouse. \**P* < 0.05. Exact *P* values shown when 0.1 > *P* > 0.05.

**Supplemental Fig. S3.**
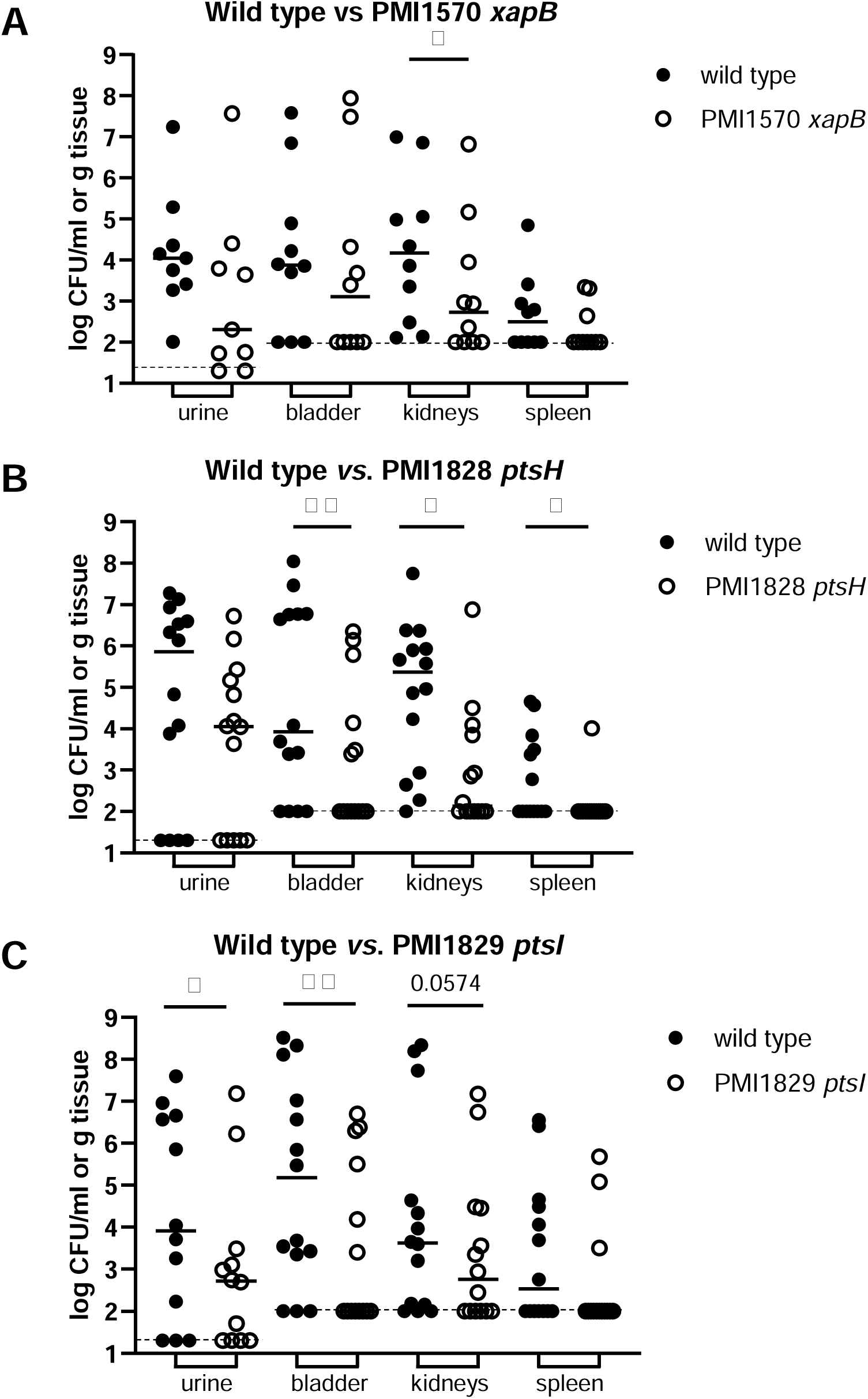
Wild type and mutant CFU recovered from 7 d murine 1:1 co-challenges shown in Fig. 2. **A,** Wild type *vs.* PMI1570 *xapB* (n = 10). **B**, Wild type *vs.* PMI1828 *ptsH* (n = 15). **C**, Wild type *vs.* PMI1829 *ptsI*. Dashed lines indicate limit of detection (urine, 20 CFU; organs, 100 CFU). \**P* < 0.05; \*\**P* < 0.01; exact values shown for 0.1 > *P* > 0.05, Mann-Whitney U test.

**Supplemental Fig. S4.**
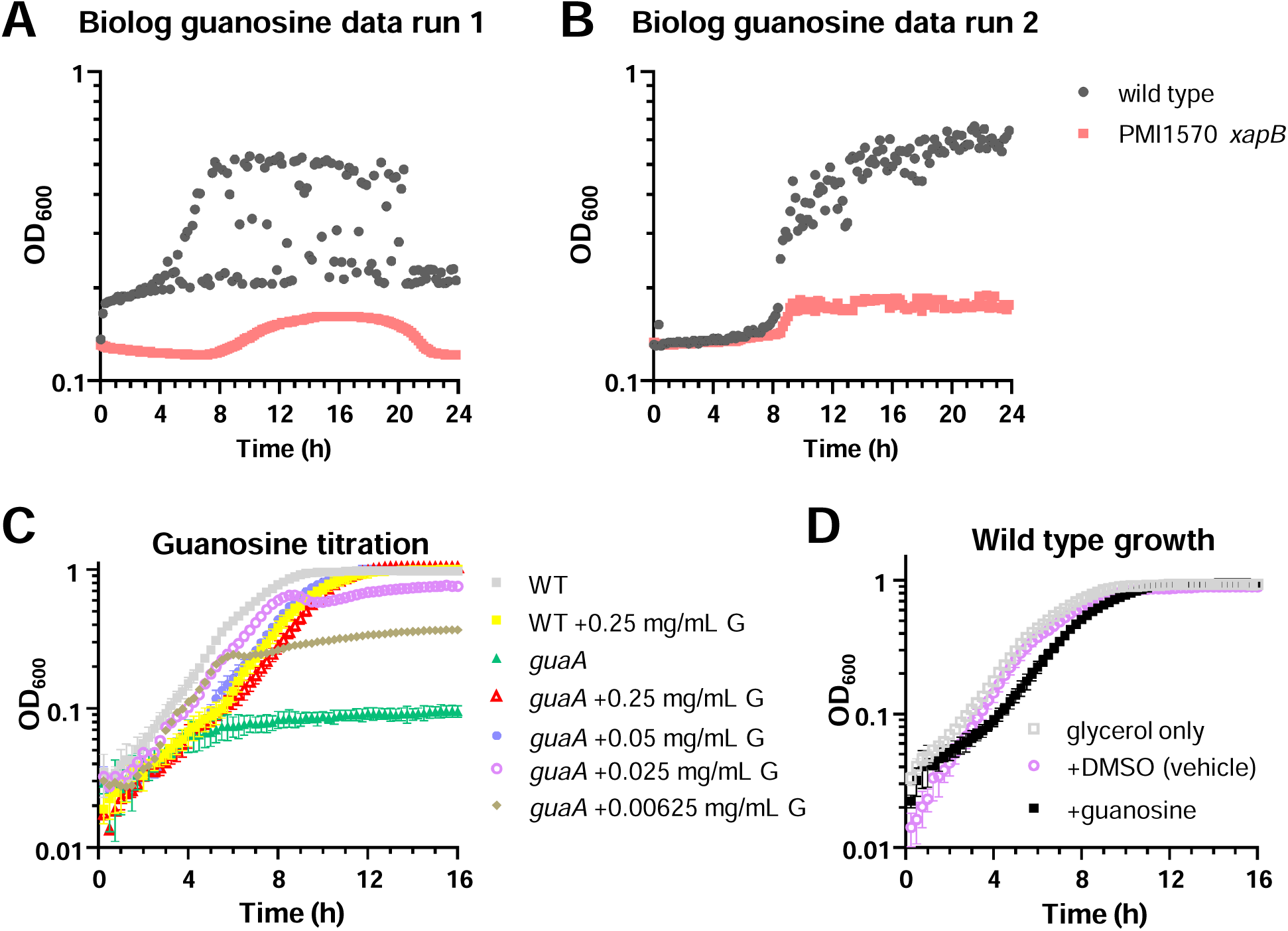
Testing guanosine as a nutrient source. **A-B**, Biolog plate PM3 suggested differential growth of wild type and PMI1570 *xapB* on guanosine as sole nitrogen source. Two independent replicates showed an erratic increase in OD_600_ for wild type but not *xapB.* **C-D**, Growth in Minimal A containing 0.2% glycerol. **C**, Titration of guanosine (G) to restore growth of the *guaA* mutant (n = 1-3). **D**, 0.05 mg/mL guanosine caused slower growth and the effect was not due to the DMSO solvent (n = 3). Error bars show SD.

**Supplemental Fig. S5.**
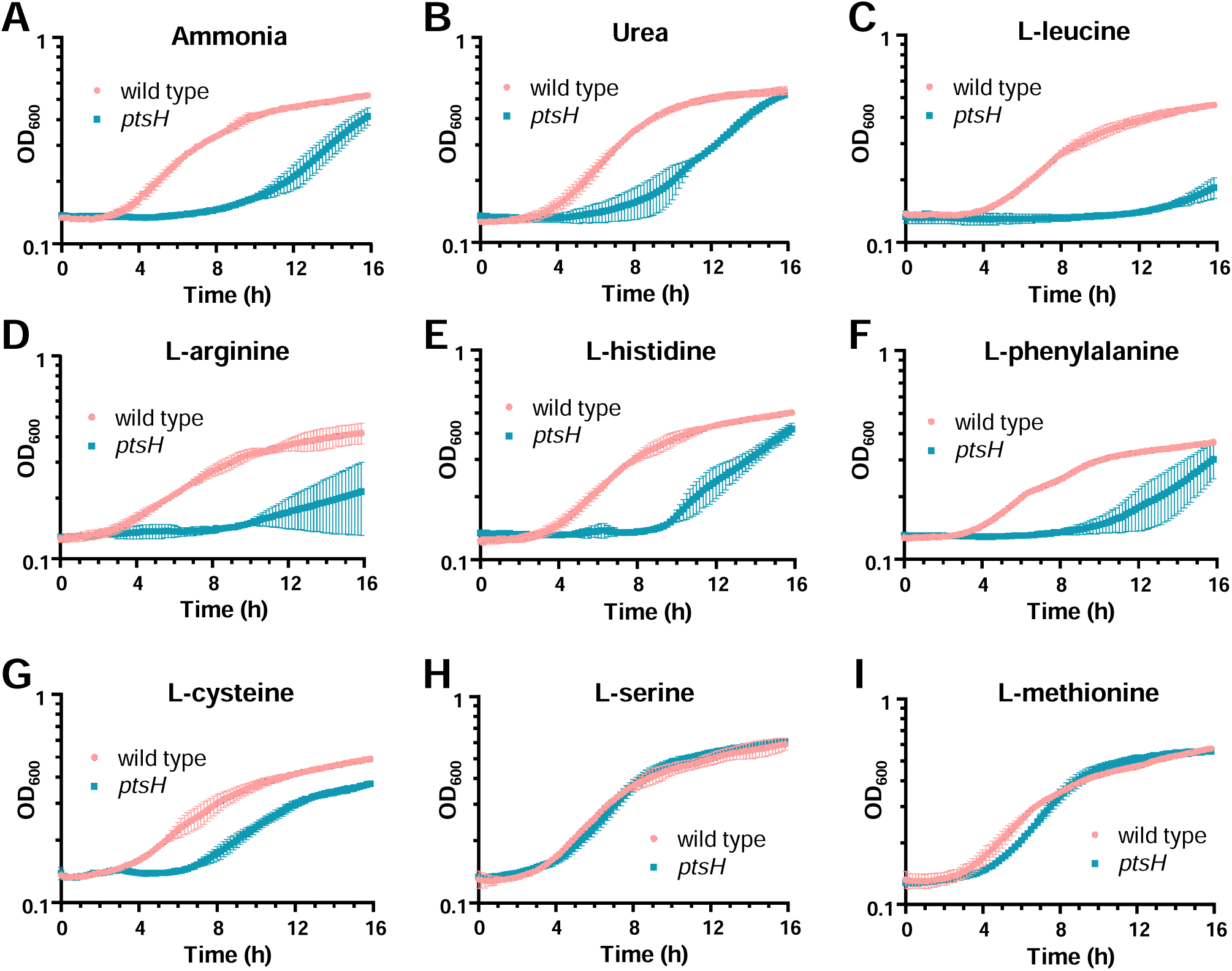
Comparison of wild type *vs. ptsH* growth on selected nitrogen sources (n = 2; error bars = SD). **A-B**, growth defect for *ptsH* mutant using preferred nitrogen sources. **C-G**, growth defects for *ptsH* mutant on selected amino acid nitrogen sources. **H-I**, not all amino acids produced reduced growth for the *ptsH* mutant.

**Supplemental Fig. S6.**
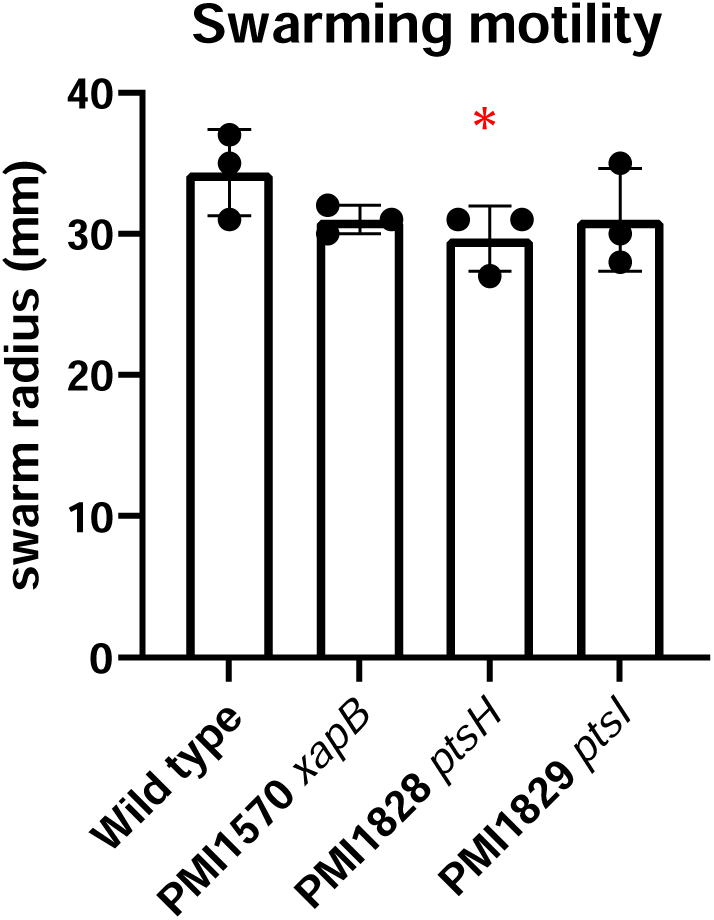
Swarming motility of transporter mutants. PMI1828 *ptsH* mutant had a modest but significant decrease in swarming motility (n = 3; error bars show SD). \**P* < 0.05, one-way ANOVA *vs.* wild type with Dunnett’s multiple comparisons test.

**Supplemental Fig. S7.**
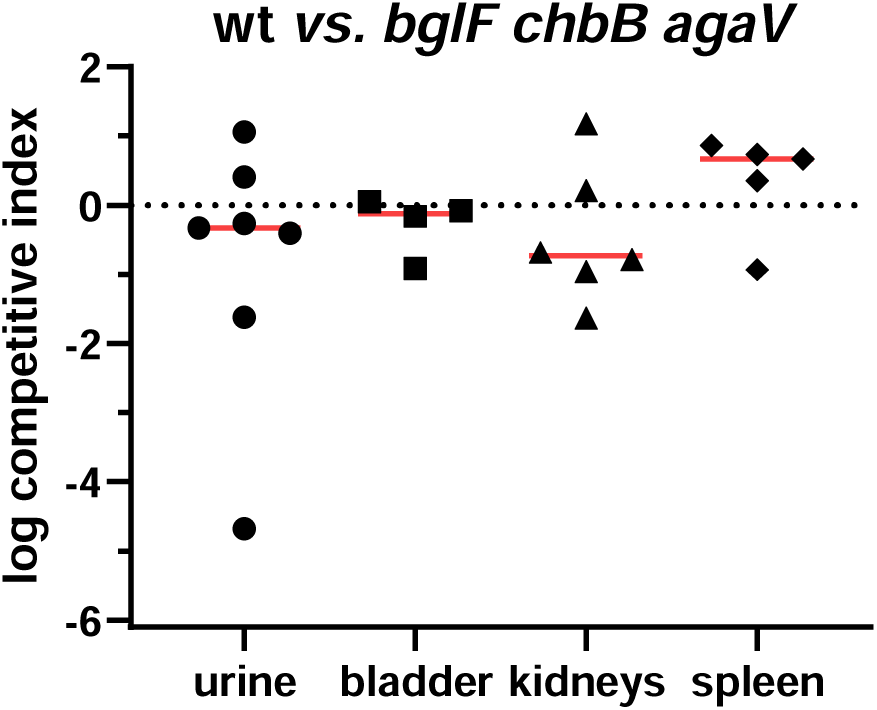
Triple mutant co-challenge of *in vivo* uninduced PTS transporters. A triple mutant with the least-induced *in vivo* PTS transporter genes had no defect in co-challenge competition with wild type. Horizontal lines show medians. Dashed line indicates equal fitness of wild type and mutant (log CI= 0).

**Supplemental Fig. S8.**
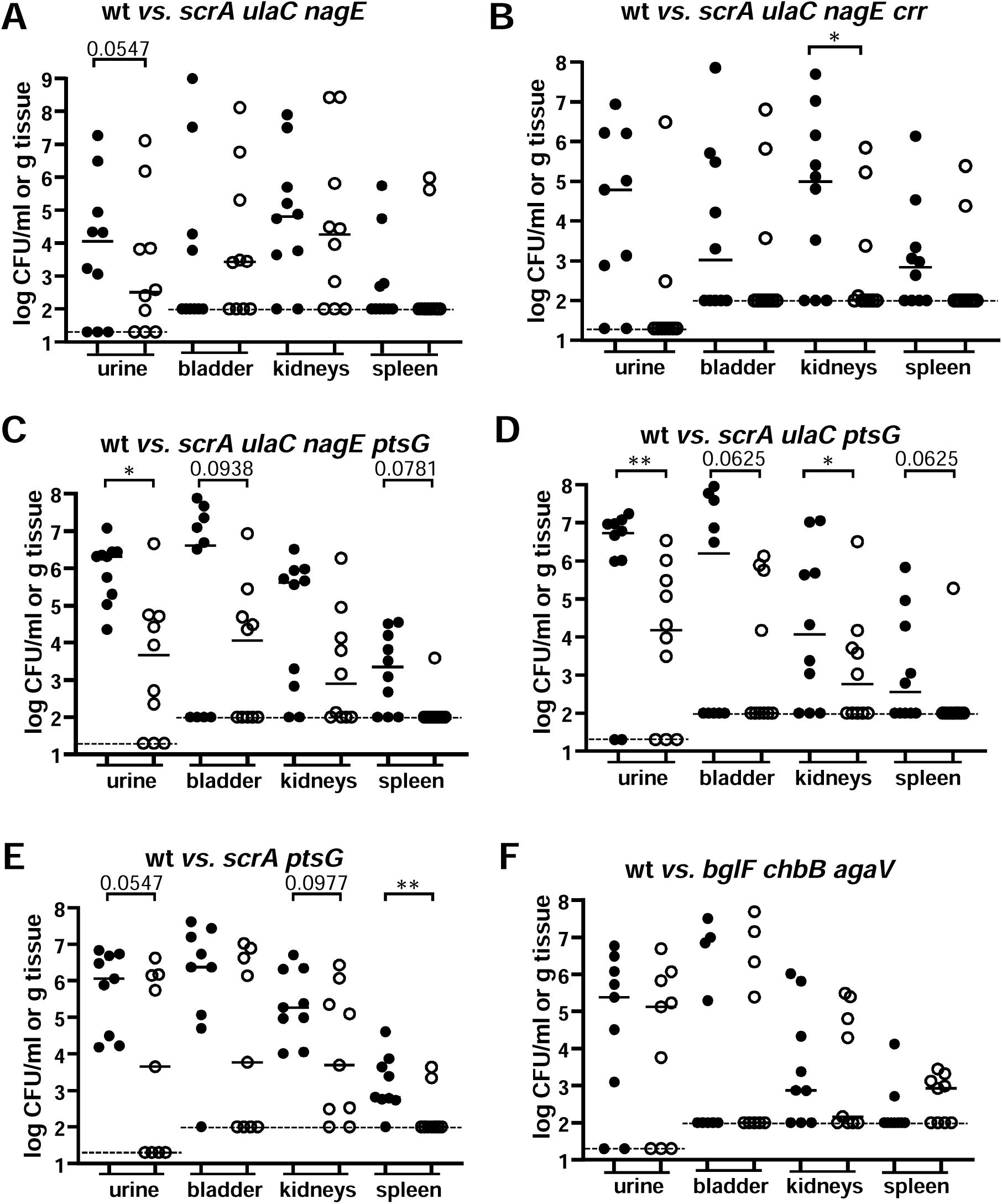
PTS multi-mutant co-challenge CFU data. **A-F,** Wild type and mutant CFU recovered from 7 d murine 1:1 co-challenges shown in Fig. 7. Solid circles are wild type (wt) and open circles are mutant (n = 10 per co-challenge). Dashed lines indicate limit of detection (urine, 20 CFU; organs, 100 CFU). \**P* < 0.05; \*\**P* < 0.01; exact values shown for 0.1 > *P* > 0.05, Mann-Whitney U test.

**Supplemental Fig. S9.**
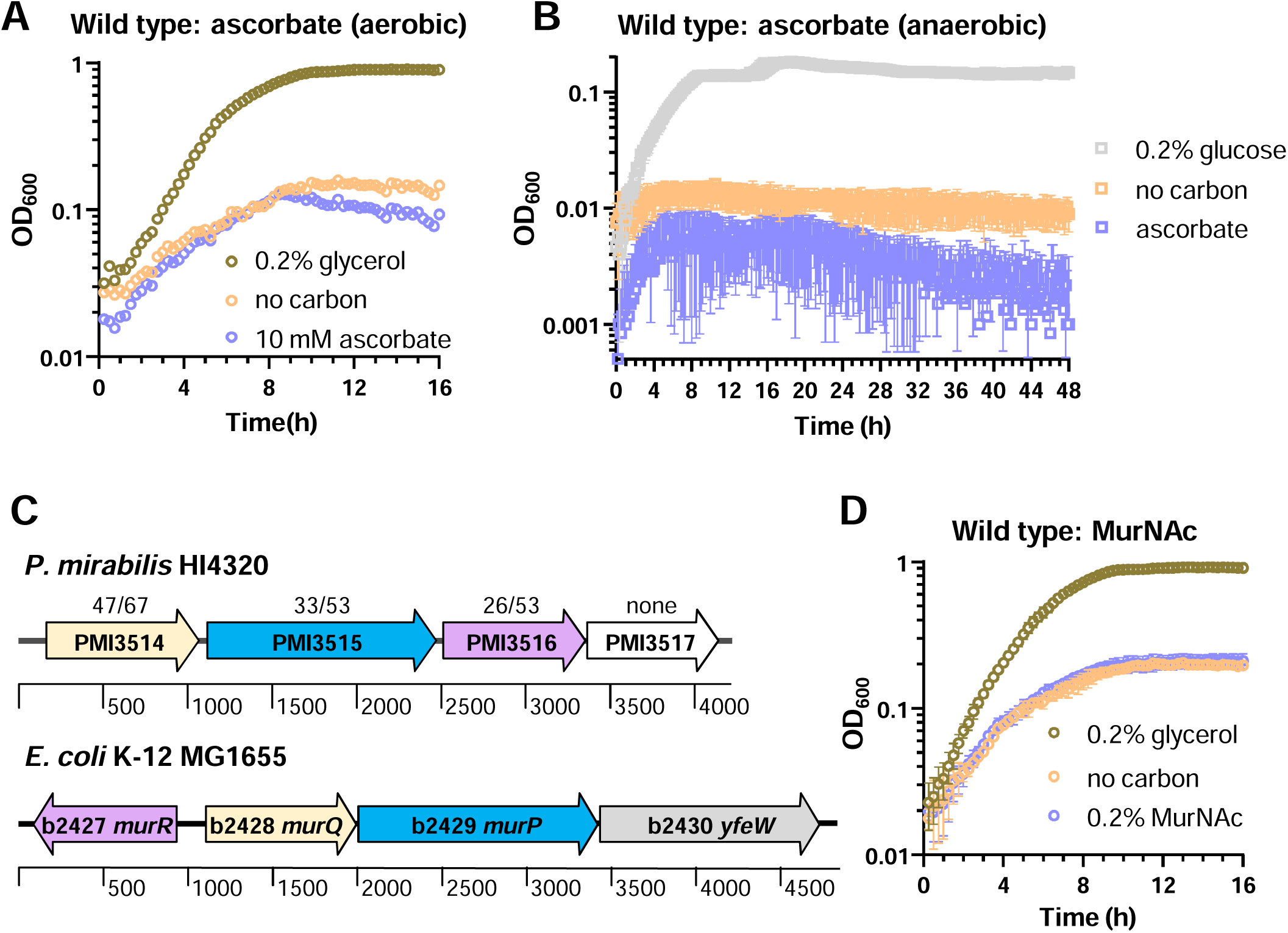
Substrates remain unconfirmed for both UlaC and ScrA. **A-B**, ascorbate growth curves with wt *P. mirabilis* HI4320. **A**, aerobic atmosphere (n = 1). **B,** anaerobic atmosphere (n = 2). In B, glucose was used as the carbon source for the positive control. **C-D,** further investigation of the PMI3515 (*scrA*) locus and substrate. **C,** Organization of PMI3515 transporter locus compared with *E. coli mur* locus. Colors indicate genes encoding proteins with similar functions, and numbers above genes indicate % identity/similarity with the same-colored predicted protein. **D,** Growth curves in Minimal A. Using MurNAc as the sole carbon source did not allow growth by wild-type HI4320 (n = 2). Error bars = SD.

**Supplemental Fig. S10.**
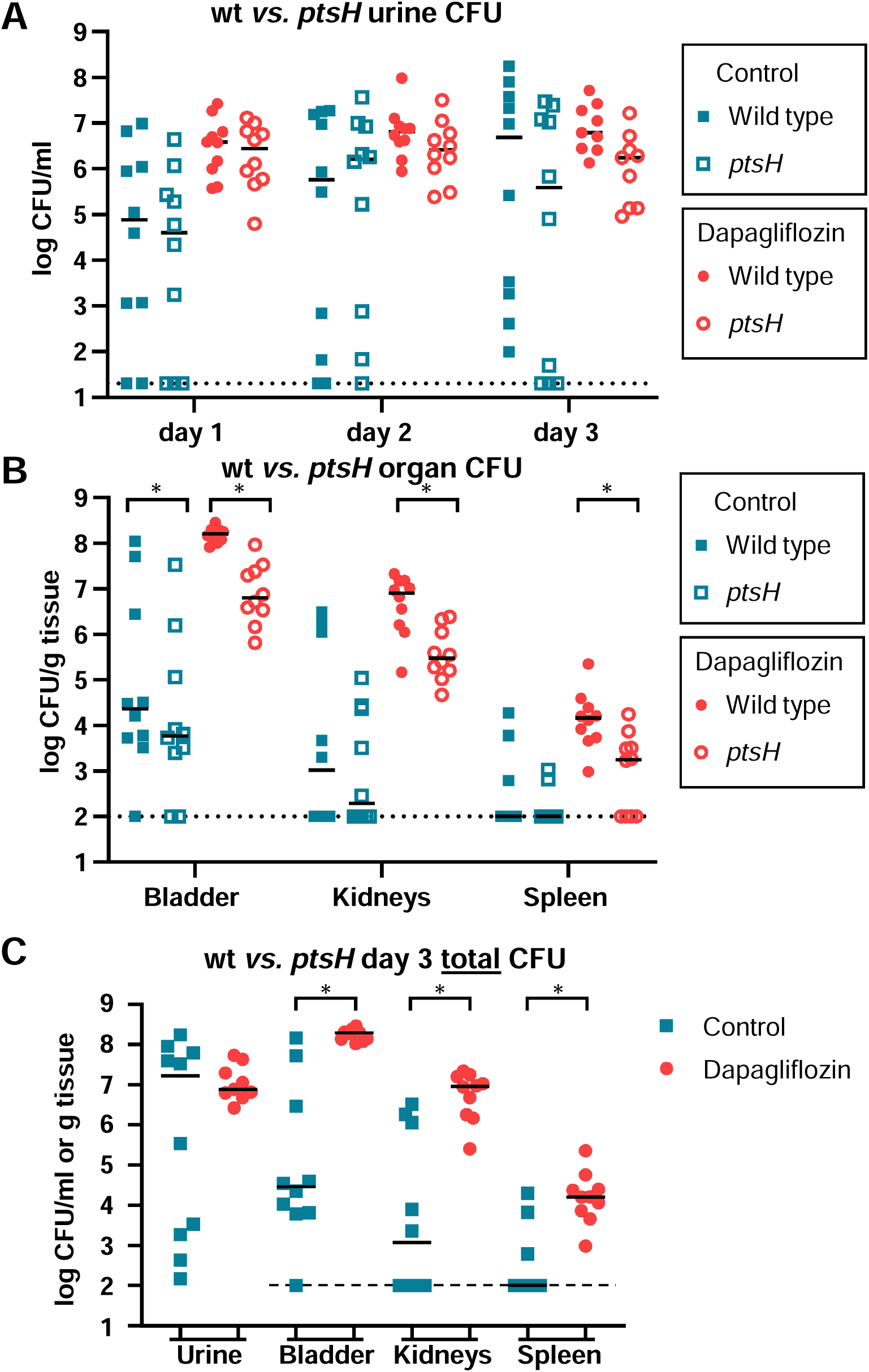
Bacterial recovery from mice co-challenged with 1:1 wild type *vs. ptsH* mutant. Mice were either treated with dapagliflozin or received normal water (control). Horizontal lines indicate medians. **A,** urine CFU at days 1, 2, or 3 post-inoculation. **B**, bacterial recovery from tissues 3 d post-inoculation. A-B, \**P*_ad_j = below threshold, multiple Wilcoxon tests with Holm-Šídák correction. **C**, Total CFU (wt + *ptsH*) shows higher overall colonization during hyperglucosuria. \**P*_ad_j = below threshold, multiple Mann-Whitney tests with Holm-Šídák correction. **A-C**, Dashed lines indicate limit of detection (urine = 20; organs = 100).

### Supplemental Datasets

**Supplemental Dataset S1. Group 1 In-seq EdgeR outputs.** Sequencing reads containing the end of the targetron insertion were aligned with the *P. mirabilis* HI4320 genome and analyzed for relative mutant recovery using EdgeR. The spreadsheet tabs are 1) overall statistics for each mutant in Group 1; 2) urine *vs.* input; 3) kidney *vs.* input; 4) bladder *vs.* input; 5) input spiral (control for outgrowth on agar) *vs.* input; and 6) list of abbreviations and nomenclature.

**Supplemental Dataset S2. Group 2 In-seq EdgeR outputs.** Sequencing reads containing the end of the targetron insertion were aligned with the *P. mirabilis* HI4320 genome and analyzed for relative mutant recovery using EdgeR. The spreadsheet tabs are 1) overall statistics for each mutant in Group 2; 2) urine *vs.* input; 3) kidney *vs.* input; 4) bladder *vs.* input; 5) input spiral (control for outgrowth on agar) *vs.* input; and 6) list of abbreviations and nomenclature.

